# Developmental cognitive neuroscience using Latent Change Score models: A tutorial and applications

**DOI:** 10.1101/110429

**Authors:** Rogier A. Kievit, Andreas M. Brandmaier, Gabriel Ziegler, Anne-Laura van Harmelen, Susanne M. M. de Mooij, Michael Moutoussis, Ian Goodyer, Ed Bullmore, Peter B. Jones, Peter Fonagy, the NSPN Consortium, Ulman Lindenberger, Raymond J. Dolan

## Abstract

Assessing and analysing individual differences in change over time is of central scientific importance to developmental neuroscience. However, the literature is based largely on cross-sectional comparisons, which reflect a variety of influences and cannot directly represent change. We advocate using *latent change score* (LCS) models in longitudinal samples as a statistical framework to tease apart the complex processes underlying lifespan development in brain and behaviour using longitudinal data. LCS models provide a flexible framework that naturally accommodates key developmental questions as model parameters and can even be used, with some limitations, in cases with only two measurement occasions. We illustrate the use of LCS models with two empirical examples. In a lifespan cognitive training study (COGITO, N=204 (N=32 imaging) on two waves) we observe correlated change in brain and behaviour in the context of a high-intensity training intervention. In an adolescent development cohort (NSPN, N=176, two waves) we find greater variability in cortical thinning in males than in females. To facilitate the adoption of LCS by the developmental community, we provide analysis code that can be adapted by other researchers and basic primers in two freely available SEM software packages (lavaan and Ωnyx).

**Highlights:**

- We describe Latent change score modelling as a flexible statistical tool
- Key developmental questions can be readily formalized using LCS models
- We provide accessible open source code and software examples to fit LCS models
- White matter structural change is negatively correlated with processing speed gains
- Frontal lobe thinning in adolescence is more variable in males than females

## 1 Introduction

*When thinking about any repeated measures analysis it is best to ask first, what is your model for change? (McArdle, 2009, p. 579)*

Developmental cognitive neuroscience is concerned with how cognitive and neural processes change during development, and how they interact to give rise to a rich and more or less rapidly fluctuating profile of cognitive, emotional and behavioural changes. Many, if not all, central questions in the field can be conceived as related to the temporal dynamics of multivariate brain-behaviour relations. Theories in developmental cognitive neuroscience often implicitly or explicitly suggest causal hypotheses about the direction of the association between variables of interest, the temporal precedence of their emergence, and the likely consequences of interventions. For instance, the *maturational viewpoint* (e.g. Gesell, 1929; cf. Johnson, 2011; Segalowitz & Rose-Krasnor, 1992) proposes that development of key brain regions (e.g. the frontal lobes) is a necessary precondition to acquiring psychological capacities (e.g. cognitive control or inhibition). This represents a clear causal pathway, where developmental change in neural regions precedes, and causes, changes in faculties associated with those regions (also known as *developmental epigenesis*). This is contrasted with *interactive specialisation theory* (Johnson, 2011), where *probabilistic epigenesis* posits bidirectional causal influences from mental function to brain structure and function. These competing theories make explicit claims about the temporal order of development, as well as the causal interactions between explanatory levels. Similarly, *developmental mismatch theory* (Ahmed, Bittencourt-Hewitt, & Sebastian, 2015; Mills, Goddings, Clasen, Giedd, & Blakemore, 2014; Steinberg, 2008; van den Bos & Eppinger, 2016) suggests that a key explanation of risk taking behaviour in adolescence is the delayed development of brain regions associated with cognitive control (e.g. the frontal lobe) compared to regions associated with mediating emotional responses (e.g. the amygdala). This too posits a clear brain-behaviour dynamic, where a mismatch between maturation in executive brain regions compared to emotion systems is hypothesized to affect the likelihood of certain (mal)adaptive behaviours. Empirical examples show, for instance, that frontoparietal structural connectivity (but not functional connectivity) determined longitudinal changes in reasoning ability (Wendelken et al., 2017).

An active area of research where cognitive or behavioural changes are presumed to precede changes in brain structure or function is that of *training-induced plasticity*. For instance, Bengtsson et al. (2005) found that degree and intensity of piano practice in childhood and adolescence correlated with regionally specific differences in white matter structure, and that this effect was more pronounced in developmental windows in which maturation was ongoing. This was interpreted as evidence of training-induced plasticity, suggesting that behavioural modifications (i.e. prolonged practice) preceded, and caused, measurable changes in white matter structure^1^. More direct longitudinal evidence in 845 children scanned on two occasions suggests that grey matter volume and changes in white matter microstructure are slower in individuals with more severe psychiatric symptoms, but not vice versa. This is compatible with (although not conclusive evidence of) the hypothesis that differences in structural changes are consequences, not causes, of psychiatric symptoms (Muetzel et al., 2017).

Although such questions are characterized by a fundamental interest in temporal dynamics and causality, much of the literature is dominated by cross-sectional (age-heterogeneous) data that are ill equipped to resolve these questions (Lindenberger & Poetter, 1998; Lindenberger, von Oertzen, Ghisletta, & Hertzog, 2011; Salthouse, 2014). For instance, individual differences in brain structure may precede differential changes in cognitive abilities (e.g. certain clinical conditions), or changes in cognitive abilities may trigger measurable changes in brain structure (e.g. learning-induced plasticity). Although these hypotheses imply radically different causal pathways and (potential) intervention strategies, they are often indistinguishable in cross-sectional data. Moreover, aggregated cross-sectional data can be affected by cohort effects (i.e. different populations) which in turn can lead to overestimates (e.g. cohort differences, Sliwinski, Hoffman, & Hofer, 2010), underestimates (e.g. selective attrition, training effects; Willis & Schaie, 1986), and even full reversals of the direction of effects observed between groups compared to within groups (Kievit, Frankenhuis, Waldorp, & Borsboom, 2013). Most crucially, cross-sectional aggregations do capture change at the individual level, nor individual differences in intra-individual change (Baltes, Reese, & Nesselroade, 1977). Thus, they fail to directly address the most fundamental questions of developmental science: How and why do people differ in the way they develop?

The recent rapid increase in the study of large, longitudinal, imaging cohorts (Poldrack & Gorgolewski, 2014) provides unprecedented opportunities to study these key questions. Here we introduce a class of Structural Equation Models called *Latent Change Score Models* that are specifically tailored to overcome various weaknesses of more traditional approaches, and are well suited to address hypotheses about temporal, interactive dynamics over time.

## 2 Towards a model-based longitudinal Developmental Cognitive Neuroscience

Structural equation modelling (SEM) combines the strengths of path modelling and latent variable modelling and has a long tradition in the social sciences (Bollen, 1989; Tomarken & Waller, 2005). Path modelling (an extension of (multiple) regression) allows for simultaneous estimation of multiple hypothesized relationships, specification of directed relations that correspond to hypothesised causal pathways, and models in which constructs may function as both dependent and independent variables. Latent variable modelling allows researchers to use observed (manifest) variables to infer and test theories about unobserved (latent) variables.

In offering a flexible framework for multivariate analyses SEM has several key strengths compared to other methods of analysis (Rodgers & Lee, 2010). First, SEM forces researchers to posit an explicit model, representing some hypothesized explanatory account of the data, which is then compared to the observed data (usually a covariance matrix, or a covariance matrix and a vector of means). The extent to which the hypothesized model can reproduce the observations is adduced as evidence in favour of, or against, some proposed model of the construct under investigation. Moreover, SEM helps make researchers aware of assumptions that may be hidden in other approaches (e.g., assumptions of equal variances across groups)^2^. Second, by using latent variables researchers can account for measurement error in observed scores. This strategy not only increases power to detect true effects (van der Sluis, Verhage, Posthuma, & Dolan, 2010), but also offers greater validity and generalizability in research designs (Little, Lindenberger, & Nesselroade, 1999). Specifically, it can be used to test for bias across subgroups (e.g. tests functioning differently for different subgroups, Wicherts, Dolan, & Hessen, 2005) and biased estimates across developmental time (Wicherts et al., 2004), and improve the use of covariates (Westfall & Yarkoni, 2016).

In recent decades, various extensions of SEM have been developed for longitudinal, or repeated measures, data (McArdle, 2009). More traditional techniques (e.g. repeated measures ANOVA) are rarely tailored to the complex error structure of longitudinal data, neglect individual differences and are not developed explicitly to test the predictions that follow from causal hypotheses across a whole set of variables simultaneously. The longitudinal SEM framework, closely related to general linear mixed modelling (Bernal-Rusiel et al., 2013; Rovine & Molenaar, 2001), is so flexible that many common statistical procedures such as t-tests, regressions, and repeated measures (M)ANOVA can be considered special cases of longitudinal SEM models (Voelkle, 2007). Common procedures in developmental cognitive (neuro)science including cross-lagged panel models or simple regressions (on either raw or difference scores) can be considered special cases of LCSM’s, but without various benefits associated with SEM such as reduction of measurement error and incorporation of stable individual differences (Hamaker, Kuiper, & Grasman, 2015).

Examples of longitudinal SEM include latent growth curve models, latent change score models, growth mixture models, latent class growth curve modelling and continuous time modelling (Driver, Oud, & Voelkle, 2016; McArdle, 2009). In the next section we describe a specific subtype of longitudinal SEM known as the *Latent Change Score Models (LCS,* sometimes also called *Latent Difference Score* models) (McArdle & Hamagami, 2001b; McArdle & Nesselroade, 1994). This particular class of models is especially versatile and useful for researchers in developmental cognitive neuroscience as it can model change at the construct level, can be used with a relatively modest number of time points (a minimum of 2, although more are desirable) and is especially powerful for testing cross-domain (i.e. brain behaviour) couplings.

## 3 The Latent Change Score model

Latent Change Score models (McArdle & Hamagami, 2001b; McArdle & Nesselroade, 1994) are a powerful and flexible class of Structural Equation Models that offer ways to test a wide range of developmental hypotheses with relative ease. LCSMs have been used to considerable effect in developmental (cognitive) psychology to show a range of effects including that vocabulary affects reading comprehension but not vice versa (Quinn, Wagner, Petscher, & Lopez, 2015), that people with dyslexia show fewer intellectual benefits from reading than controls (Ferrer, Shaywitz, Holahan, Marchione, & Shaywitz, 2010), that positive transfer of cognitive training generalizes beyond the item-level to cognitive ability (Schmiedek, Lövdén, & Lindenberger, 2010), that volume changes of the hippocampus and prefrontal white matter are reliably correlated in adulthood and old age (Raz et al., 2005), that an age-related decline in white matter changes is associated with declines in fluid intelligence (Ritchie et al., 2015) and that basic cognitive abilities such as reasoning and vocabulary show mutualistic benefits over time that may partially explain positive correlations among cognitive abilities (Kievit et al., 2017). One of the first applications of LCS in cognitive neuroscience showed that ventricle size in an elderly population predicted rate of decline on memory tests across a seven year interval (McArdle et al., 2004). There are several excellent tutorials on longitudinal SEM (Ghisletta & McArdle, 2012; Grimm, 2007; Jajodia & Archana, 2012; McArdle & Grimm, 2010; Petscher, Quinn, & Wagner, 2016; Snitz et al., 2015; Usami, Hayes, & McArdle, 2016; Zhang, Hamagami, Grimm, & McArdle, 2015), and the approach we outline below builds heavily upon this previous work, whereas we illustrate LCS models specifically in the context of Developmental Cognitive Neuroscience.

We will start with the simplest model, using one variable measured on two occasions, and then present four extensions of the model. These extensions will sequentially incorporate latent variables, add multiple coupled domains (cognitive and neural measures), extend to multiple time waves (latent growth- and dual change score models) and finally test for differences in multiple groups. After discussing each of the basic models in turn, we will cover methodological challenges including estimation, model fit, interpretation and model comparison. For each of the five types of models we discussed below, we provide example syntax that simulates data under a selected parameterisation and fits the model in question to the data. These scripts are freely available at the Open Science Framework ^https://osf.io/4bpmq/?view_only=5b07ead0ef5147b4af2261cb864eca32^ to be used, modified and extended by the wider community.

### 3.1 Univariate Latent Change Score model

Imagine a researcher studying a psychological variable of interest, repeatedly measured at two time points (T1 and T2) in a population of interest. A traditional way to examine whether scores of a group of individuals increased or decreased between T1 and T2 is performed by means of a paired t-test. Using some simple modifications, the LCS allows us to go beyond this traditional analysis framework even in this simplest case. The basic steps of a univariate latent change score model are as follows. To facilitate understanding, in the examples below, we will use informative notation (e.g. ‘COG’ for cognitive measures and ‘NEU’ for neural measures). For a more standard mathematical notation of the LCS we refer the reader to texts such as (McArdle, 2008; McArdle & Hamagami, 2001a; Newsom, 2015; Petscher et al., 2016). First, we conceptualize the scores of an individual *i* on the construct of interest *COG* at time *t* as being a function of an autoregressive component and some residual. By fixing the regression weight of *COGT2* on *COGT1* to 1, the autoregressive equation simplifies to

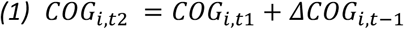

From this it follows that the change score is simply:

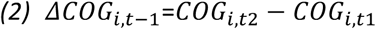

The powerful step in the context of SEM is to define a latent change score factor *ΔCOG1*, which is measured by time point 2 with a factor loading fixed to 1. Doing so creates a latent factor that captures the change between time 1 and time 2. Finally, we can add an regression parameter *β* to the change score, which allows us to investigate whether the degree of change depends on the scores at time 1 as follows:

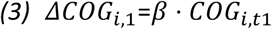

With this model in place we can address three fundamental questions. The first and simplest question is whether there is a reliable average change from T1 to T2. This is captured by the mean of the latent change factor, *μΔCOGT1*. Under relatively simple assumptions this test is equivalent to a paired t-test (Coman et al., 2013). However, even this simplest implementation of the latent change score model yields two additional parameters of considerable interest. First, we can now estimate the *variance* in the change factor, *σ^2^ΔCOG1*, which captures the extent to which individuals *differ* in the change they manifest over time. Second, we can specify either a covariance or an autoregressive parameter *β* which captures the extent to which change is dependent, or *proportional,* to the scores at time one (this parameter can also be specified as a covariance if so desired). Note that if a regression parameter is included, the mean change should be interpreted conditional on the regression path – A covariance will yield the mean ‘raw’ change.

SEMs are often illustrated using *path models*. Such representations go back to Wright (1920), and allow researchers to represent complex causal patterns using simple visual representations. Figure 1 shows the commonly employed symbols, meaning and notation. The simplest representation of the univariate latent change model is shown in Figure 2, and can be fit to data measured on two occasions. As the autoregressive parameter between *COGT2* and *COGT1* is fixed to unity, we implicitly assume that the intervals are equidistant across individuals. Deviations from this assumption can be dealt with by rescaling scores (Ferrer & McArdle, 2004, p. 941) or, ideally, by using definition variables (Mehta & West, 2000) or continuous-time modelling approaches (Driver et al., 2016), which yield parameters that more easily generalize across different longitudinal designs. The model shown in Figure 2 is *just identified*, that is, there are as many unique pieces of information that enter the model (two variances, two means and a covariance) as parameters to be estimated (one observed variance, one latent variance, one observed mean score, one latent mean score and one regression parameter). This means that although we can estimate this model, we cannot interpret model fit in isolation unless additional data (multiple waves, multiple domains or multiple indicators) are included. However, we can make use of parameter constraints, namely fixing certain parameters to zero and employ likelihood ratio testing of hypotheses about specific parameters. For instance, one can separately fit two similar models: once a model with the latent change variance parameter freely estimated, and once with the variance constrained to 0 (implying no differences in change). The difference in model fit will be chi-square distributed with *k* degrees of freedom, where *k* is the number of parameters constrained to equality (Neale, 2000; but see Stoel, Garre, Dolan, & van den Wittenboer, 2006). If fixing the variance of change parameter to 0 leads to a significant drop in model fit, it would suggest that individuals change heterogeneously. However, it should be noted that this inference is only valid compared to the simpler model - a more extensive model with additional variables or time points may lead to different conclusions about heterogeneity in change. A more practical concern is that constraining variances parameters may lead to failure of model convergence which renders interpretation challenging – See section 4.1.2 for more guidance. Similar procedures can be employed for any other parameter of interest or combinations of any number of parameters. The likelihood ratio test is especially suitable for parameters such as variances, as the simplifying assumptions of parameter significance tests such as the often-used Wald test may not hold (i.e. a variance cannot be negative). Next, we examine how to extend the LCS model to include latent variables.

**Figure 1:**
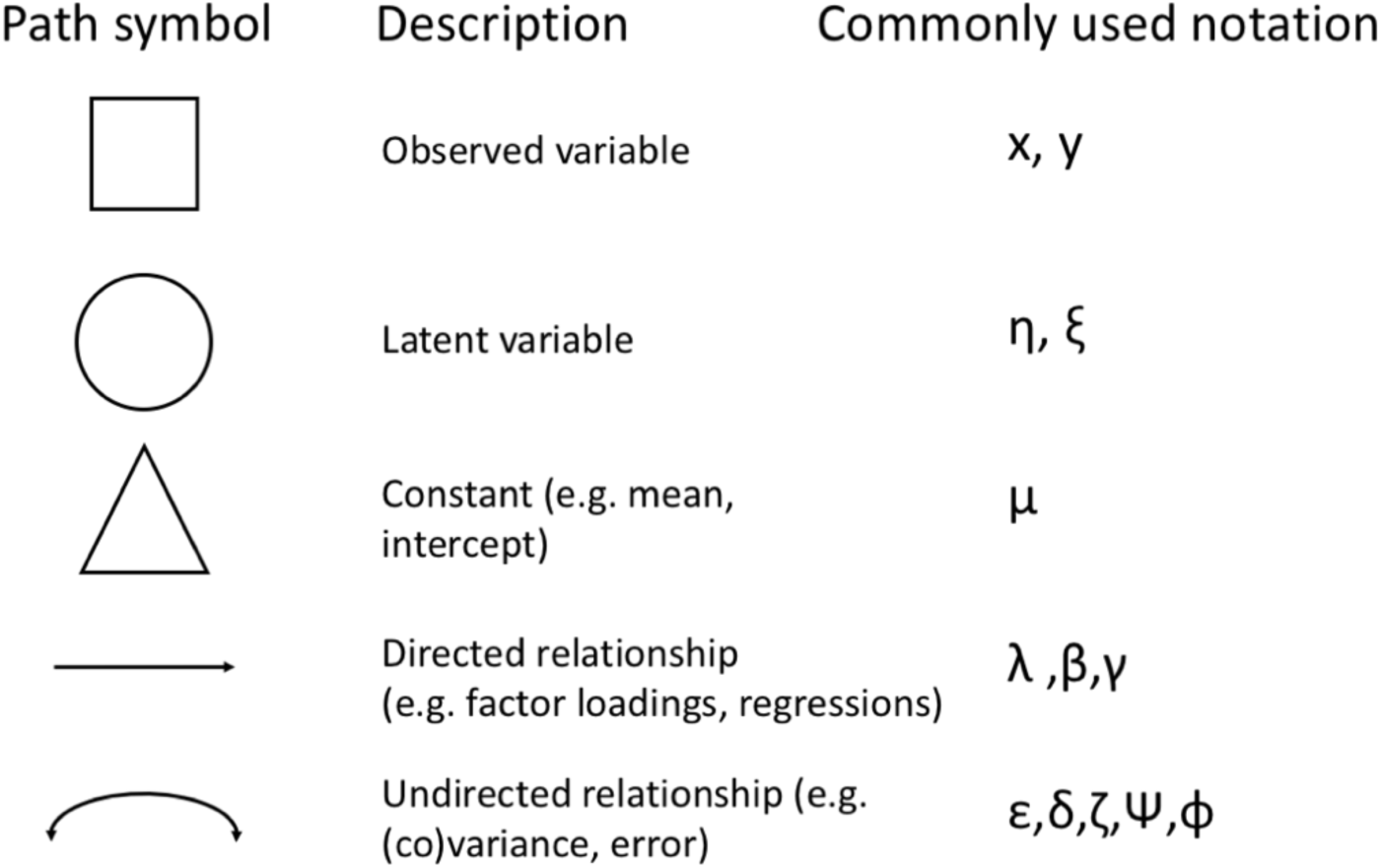
Basic path model notation

**Figure 2:**
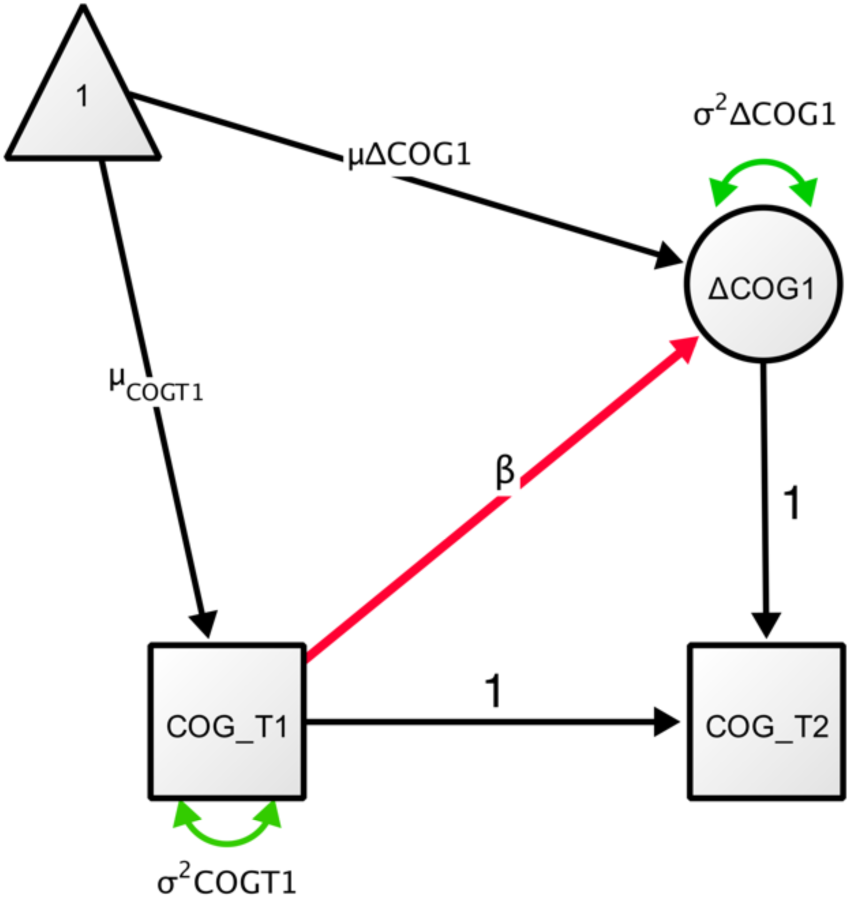
Univariate Latent Change Score Model. Variable COG is measured at two time points (COG_T1 and COG_T2). Change (DCOG1) between the two timepoints is modelled as latent variable).

### 3.2 Multiple Indicator Univariate Latent Change Score model

The above example uses a single observed variable, which was assumed to be measured without error. We can easily extend this model to have an explicit measurement model by replacing the observed score with a *latent variable*, measured by a set of observed variables. We refer to this representation as a multiple indicator latent change score model, as our aim is to model change in the latent score rather than observed scores. To do so, we model a latent variable in the manner of a traditional confirmatory factor analysis, by expressing the strength of the association between the latent variable *COG* in individuals *i* (*i*=1,.*N*) measured at times *t* (*t*=1,…*t*) and the observed scores X (*j*=1,…*j*) with factor loadings λ and error terms δ as follows:

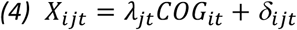

A simple multiple indicator latent change score model is shown in Figure 3. We model the mean, variance and autoregressive changes in *COG* as before, but now add a set of three (*X1-X3*) observed measurements on two occasions that are each reflections of the underlying cognitive construct of interest. Additionally, we allow for residual covariance of error terms across time points for each observed score with itself, represented as double-headed arrows. These so-called ‘correlated errors’ allow for indicator-specific variance across occasions and are generally included as default (Newsom, 2015, p. 103; Wheaton, Muthen, Alwin, & Summers, 1977). This model is similar to the univariate latent change score model in terms of the key questions it can address (rate of change *μΔCOGT1*, variance in change *σ^2^ΔCOG1*, and the relation between *COG1* and *ΔCOG1* captured by *β*), but adds the benefits of removing measurement error and establishing measurement invariance over time and (if necessary) across groups, improving inferences.

**Figure 3:**
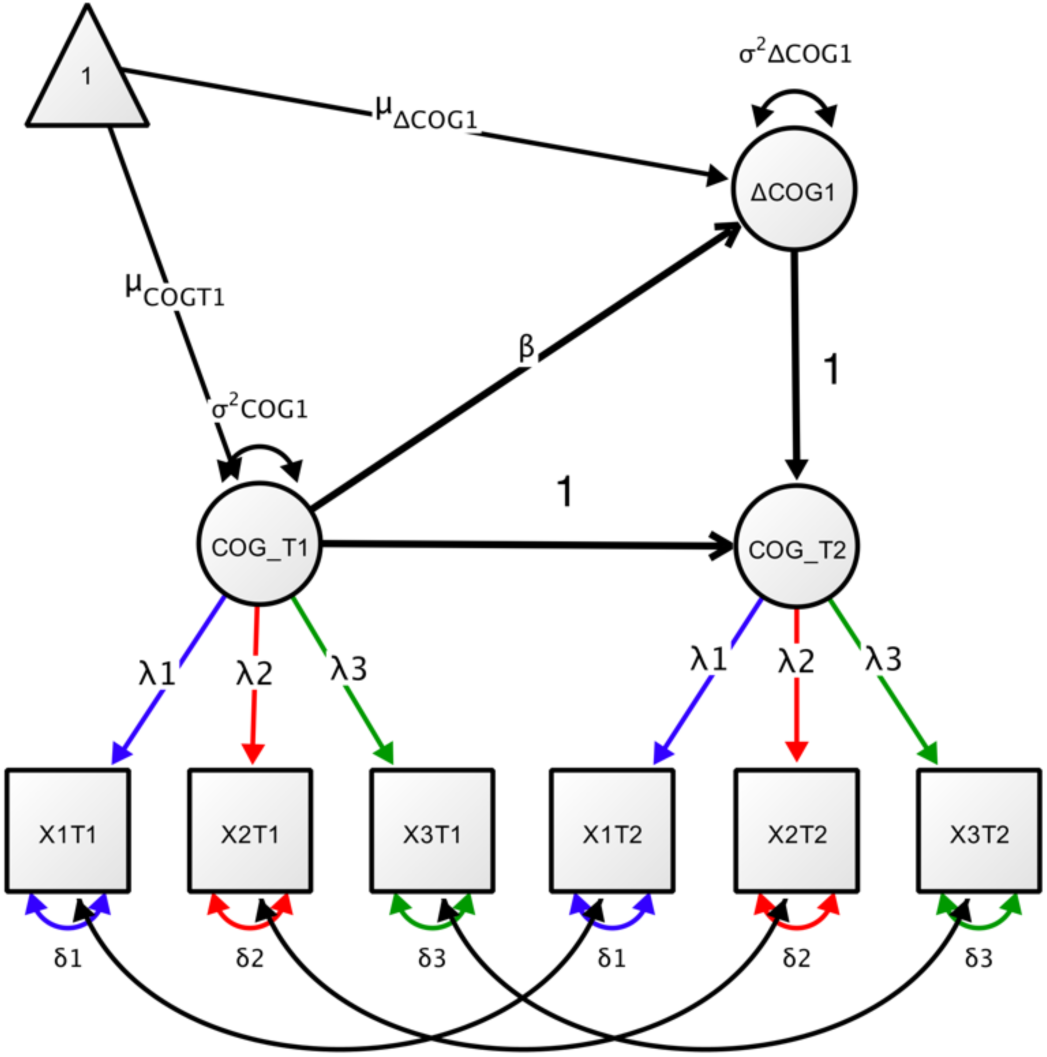
Multiple indicator univariate latent change score model. The latent construct of interest (COG) is measures at two time oints (COG_T1 and COG_T2) each measured using three indicators (X1, X2, X3). We assume measurement invariance and correlated residual errors over time. See text for a detailed description of the model parameters.

### 3.3 Bivariate Latent Change Score model

A further extension of the latent change score model is to include a second (or third, fourth, etc.) domain of interest. For convenience in notation and graphical representation we will revert back to using only observed scores, but all extensions can and - where possible should - be modelled using latent (multiple indicator) factors. We can assume the second domain is some neural measure of interest (e.g. grey matter volume in a region of interest), measured on the same number of occasions as the cognitive variable (or variables). This allows for the investigation of a powerful concept known as *cross-domain coupling* (Figure 4), that captures the extent to which *change* in one domain (e.g. *ΔCOG*) is a function of the starting level in the other (i.e. *NEU*). For instance, we can quantify the extent to which *cognitive changes* between T1 and T2 are a function of brain structure (γ2) and cognition (β1) at T1 as follows:

**Figure 4:**
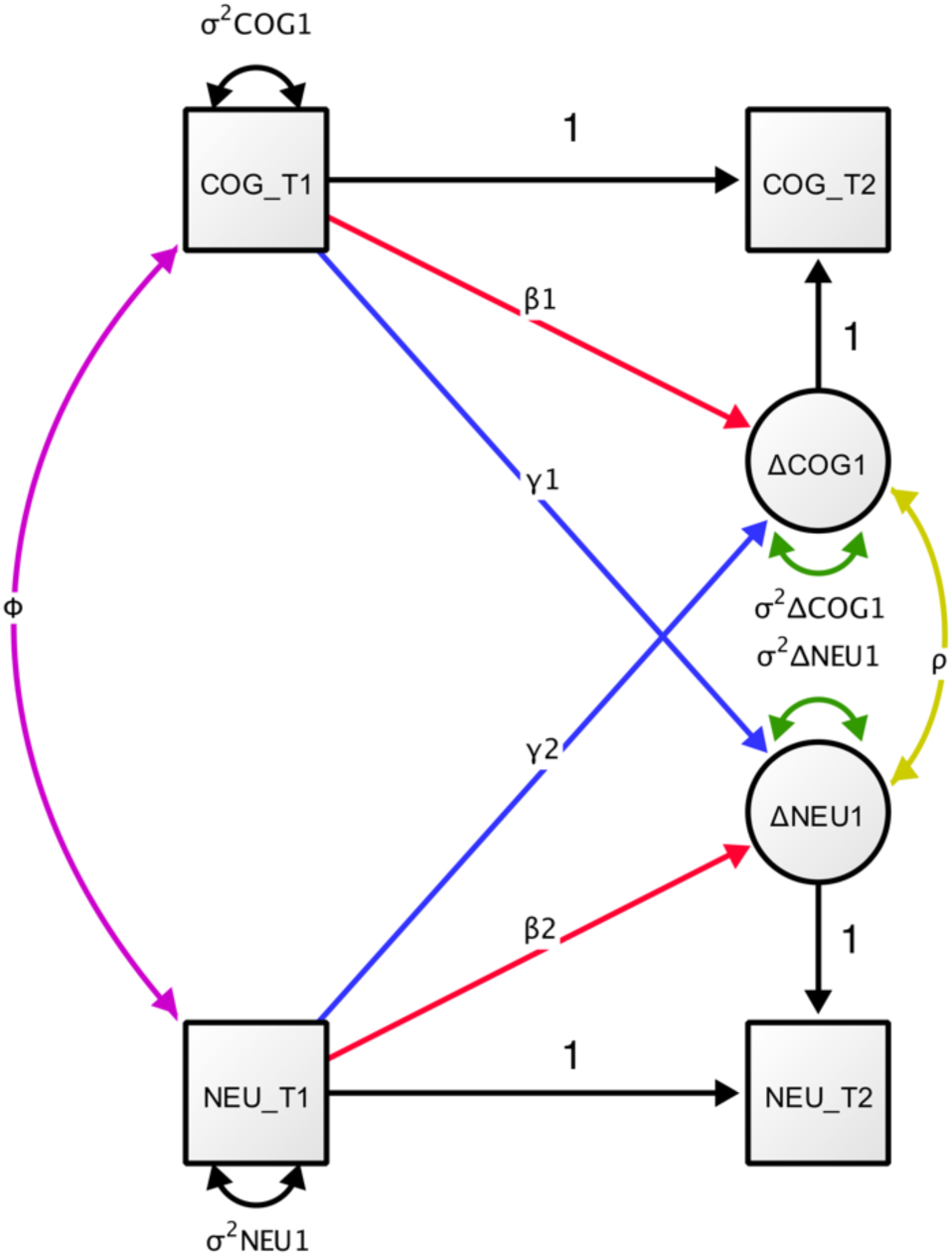
Bivariate Latent Change Score Model. Note: means are omitted for visual clarity.

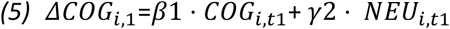

The implications for testing theories in developmental cognitive neuroscience should be immediately clear: the dynamic parameters, shown in red and blue in Figure 4, capture the extent to which changes in cognition are a function of initial condition of brain measures, vice versa or both. Likelihood ratio tests or Wald tests of these dynamic parameters (brain measures affecting rates of change in cognition, or cognitive abilities affecting neural changes) furnish evidence for, or against, models that represent uni- or bidirectional hypothesized causal influences. As is clear from Figure 4, the bivariate latent change score model can capture at least four different brain-behaviour relations of interest. First, we have brain-behaviour covariance at baseline (shown in purple), the main focus in traditional (developmental) cognitive neuroscience. Second, we have cognition to brain coupling (shown in blue, labelled γ1), where T1 scores in cognition predict the rate, or degree, of change in brain structure. For instance, the degree of childhood piano practice affected white matter structure (*ΔNEU1*) would predict a substantial cognition-to-neural coupling parameter γ1 (Bengtsson et al., 2005). Third, we have brain structure predicting rate, or degree, of cognitive change (shown in blue, labelled γ2). For example, McArdle et al. (2004) showed that ventricle size in an older population predicted rate of memory decline across an interval of 7 years. Finally, we have an estimate of correlated change (shown in yellow), reflecting the degree to which brain and behaviour changes co-occur after taking into account the coupling pathways. For instance, Gorbach et al. (2016) observed correlated change between hippocampal atrophy and episodic memory decline in older adults. More generally, correlated change may reflect a third, underlying variable influencing both domains. The bivariate latent change score provides a powerful analytic framework for testing a wide range of hypotheses in developmental cognitive neuroscience in a principled and rigorous manner.

### 3.4 Bivariate Dual Change Score model

So far, we focused on the simplest instance of longitudinal data, namely where data is measured on two occasions. This is likely to be, for the foreseeable future, the most common form of longitudinal dataset available to researchers in developmental cognitive neuroscience, and yields many benefits compared to both cross-sectional data analyses and more traditional techniques such as cross-lagged panel models or change score regression (see section 4 for more detail). However, with a greater number of timepoints, extensions within the framework of LCS models makes it easy to capture more fine-grained dynamic processes within and across domains. For instance, a sufficient number of timepoints allows one to fit what is known as a *dual change score model* (Ghisletta & Lindenberger, 2003). In this model, we specify an additional latent variable, S (for slope), that captures the global increase or decrease across all time points. This latent variable is measured by the successive change scores *ΔCOGt*, by specifying factor loadings (α) to capture a range of dynamic shapes such as linear increase or decrease. The factor loadings of the slope factor on the constant change parameter can be fixed to a priori values to capture a range of growth processes including linear (all 1) or accelerating change (e.g. 1,2,3) -– however, due to identification constraints, they cannot generally be freely estimated from the data.

The ‘dual’ aspect of this model enters by separating the global process of change captured by the slope from the more local, time point-to-time point deviations from this trajectory denoted by the self-feedback (*β*, red pathways in Figure 5) and cross-domain coupling (γ, blue pathways in Figure 5) parameters. When modelled together with a neural variable the bivariate dual change score, shown graphically in Figure 5, can be expressed as follows

**Figure 5:**
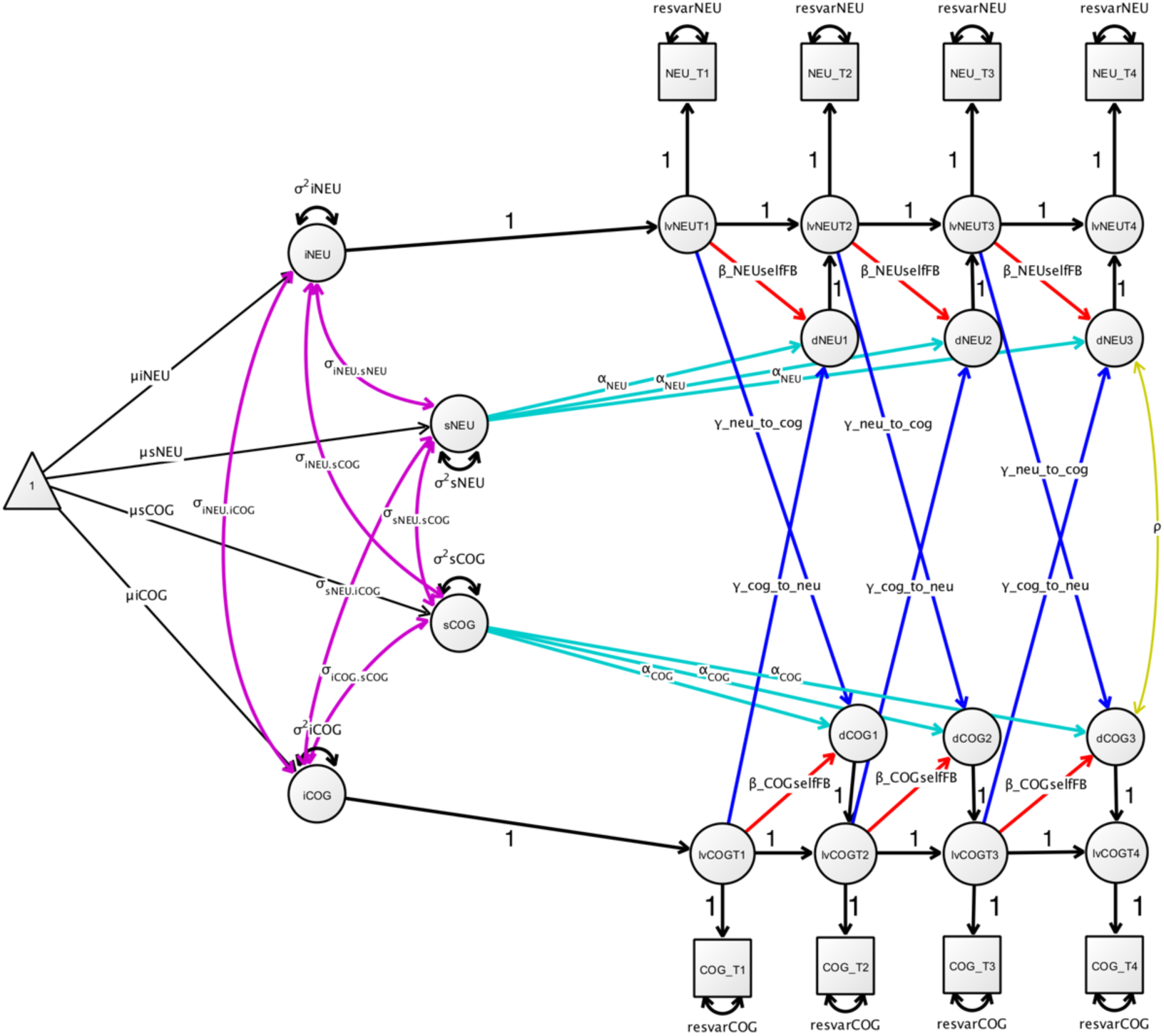
Bivariate Dual Change Score Model. This more complex latent change score model captures both the stable change over time in the form of slopes (sCOG and sNEU), as well as more fine-grained residual changes. Note this model incorporates latent variables at each timepoint – See Newsom (2015), p. 135 for more detail.

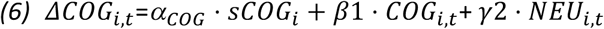

This (bivariate) dual change score model is a general approach that can capture both general trends and more fine-grained temporal dynamics. This can be especially useful when trying to separate a known, more stable change occurring during development (e.g. global cortical thinning) from more high-frequency fluctuations. The dual change score model has been used in a behaviour only context to show that (a) vocabulary influences changes in reading ability (but not vice versa) (Quinn et al., 2015); (b) and to show bivariate dynamic coupling between subjective and objective memory problems in an ageing population (Snitz et al., 2015); (c) within-person trial-to-trial RT variability predicts cognitive decline in old and very old age (Lövdén, Li, Shing, & Lindenberger, 2007); (d) perceptual speed decline in old age predicts decline in crystallized intelligence to a greater extent than vice versa (Ghisletta & Lindenberger, 2003).

### 3.5 Multigroup Latent Change Score models: Manifest groups, mixtures and intervention studies

The LCSM provides a comprehensive framework to model both within-person change across time and between-person variability in change. A final, powerful extension that can be applied to any latent change score (or SEM) model is the possibility for *multigroup* comparisons. Parameter estimates of a LCSM are valid under the assumption that there is no model misspecification and the sample is drawn from a single, homogeneous population. In practice, however, our samples may be a mixture of participants from different populations (e.g., children and older adults; men and women, low vs. high SES). There are several ways to address sample heterogeneity depending on the assumptions we are willing to make and the strength of our theoretical reasoning concerning sample heterogeneity. First and foremost, approaches to model heterogeneity can be classified by whether heterogeneity is assumed to be observed or unobserved.

When heterogeneity is observed in a confirmatory modelling approach, hypothesis testing is concerned with finding statistical evidence for difference in the key parameters of the LCSM. Cross-sectional analyses often use traditional methods such as (M)ANOVA’s to focus on simple parameters of interest, such as the mean scores on some outcome of interest. In a SEM context, it is relatively easy to compare *any* parameter of interest in a (dynamic) model across groups. To do so, one simply imposes equality constraints on the parameter of interest and compares the model where the parameter of interest is freely estimated to a constrained model as described above. Relatively sophisticated questions about changing relations between constructs, developmental dynamics and group differences can be addressed with this simple yet powerful test, with previous investigations comparing regression coefficients (e.g. the negative effect of relational bullying on friendships is stronger in boys than in girls, van Harmelen et al., 2016), or dynamic growth components (e.g. boys and girls show differential response dynamics following divorce, Malone et al., 2004). In cases where a large number of covariates are potentially relevant but we have no strong theories to guide us, more exploratory techniques such as SEM trees (Brandmaier, von Oertzen, McArdle, & Lindenberger, 2013) and SEM forests (Brandmaier, Prindle, McArdle, & Lindenberger, 2016) allows researchers to hierarchically split empirical data into homogeneous groups sharing similar data patterns, by recursively selecting optimal predictors of these differences from a potentially large set of candidates. The resulting tree structure reflects set of subgroups with distinct model parameters, where the groups are derived in a data-driven way. Group divisions are often based on observed variables, but if heterogeneity is assumed to be unobserved, researchers may turn to *latent mixture models* (McLachlan & Peel, 2005, but for a cautionary note see Bauer (2007).

An often overlooked application of LCS and SEM models is in intervention studies (McArdle, 1994). We can treat grouping of participants into treatment and control groups in precisely the same way as traditional grouping variables such as gender or education, and compare all model parameter using likelihood ratio tests. For instance, Raz et al. (2013) showed less cerebellar shrinkage in a cognitive training intervention group than in controls, and Maass et al., (2015) using SEM to demonstrate correlated change in between fitness improvement and memory. By modelling time by group interaction in a SEM context, one can use multiple indicator latent change score models to derive error-free effect sizes of the treatment effect, by subtracting average latent change in control group from average latent change in the treatment group for latent constructs (e.g. Schmiedek et al., 2010; Schmiedek, Lövdén, & Lindenberger, 2014). Once researchers have decided on which model best matches their developmental hypothesis and is compatible with the available data, it is time to estimate and interpret the model.

## 4 Challenges and limitations

### 4.1.1 Model fit, model estimation and model comparison

Once a model has been specified for a suitable dataset, a researcher will estimate the free parameters in the model. The most common approach to parameter estimation in SEM is maximum likelihood, under the assumption of multivariate normality. The extent to which this assumption is violated can bias results, although adjusted model fit indices have been developed to account for deviations from (multivariate) normality (e.g. Satorra-Bentler or Yuan-Bentler-scaled test statistics (Rosseel, 2012). Note that these methods only adjust fit indices, not standard errors, which may also be affected by deviations from (multivariate) normality – Various additional methods such as Huber-White standard errors can be used to address this challenge and are implemented in almost all SEM packages (including lavaan, see (Rosseel, 2012) p. 27 for more detail). Alternatively, other estimation strategies can be used to estimate non-continuous or non-normal outcomes (e.g., threshold models or weighted-least-squares estimators for ordinal data) but as a detailed investigation of this issue is beyond the scope of this tutorial we refer the reader to additional resources (Kline, 2011; Olsson, Foss, Troye, & Howell, 2000; Rosseel, 2012; Schermelleh-Engel, Moosbrugger, & Müller, 2003).

A key intermediate step in longitudinal SEM in the case of measurement models (e.g. Fig 3) is to provide evidence for *measurement invariance,* that is to ensure that the same latent construct (e.g. general intelligence) is measured in the same manner across time (or across groups Meredith, 1993; Millsap, 2011; Vandenberg & Lance, 2000; Wicherts et al., 2004). In other words, we want the relationship between levels of the latent variables and the observed scores to be equal across time, even when latent scores themselves are increasing or decreasing on average. Failing to establish measurement invariance can lead to incorrect conclusions about latent variables, their growth over time, and their relations to other variables (Ferrer, Balluerka, & Widaman, 2008). In longitudinal SEM, a series of increasingly strict tests (Widaman, Ferrer, & Conger, 2010) can be applied to ensure measurement invariance. Conventionally, this is done by establishing equality constraints over time, by sequentially equating the factor loadings (λ_*jt*1_ = λ_*jt*2_), error terms (δ_*jt*1_ = δ_*jt*2_) and intercepts across time or groups. Such constraints can be shown graphically in model representations – For instance in Figure 3, these equality constraints are shown by designating the same factor loading with a single label (e.g. λ1) or colour across loading (e.g. blue) across time points. Recent work shows that the CFI (comparative fit index, see for more detail below) is a practical way to test for measurement invariance across increasingly strict models created by imposing a specific sequence of model constraints (Cheung & Rensvold, 2002). If measurement invariance is violated, the extent to which inferences are affected and possible remedies using ‘partial measurement invariance’ are discussed in (Vandenberg & Lance, 2000).

A key strength of estimation in SEM is the treatment of missing data (Enders, 2001). Assuming data is either Missing Completely At Random (MCAR) or Missing At Random (MAR), which means the missing data can only be dependent on variables also measured within the same dataset (e.g. if differences in drop out are gender specific, and gender is assessed), Full Information Maximum Likelihood (FIML) can be used to estimate a model on the full dataset (including subjects with incomplete data) (Baraldi & Enders, 2010; Enders, 2001; Wothke, 2000). Using FIML for missing data (under multivariate normality) maximizes the utility of all existing data, decreases bias and increases statistical power compared to (for instance) omitting incomplete cases (‘complete case analysis’; Baraldi & Enders, 2010). In direct comparisons, FIML usually performs as well or better than alternative methods such as multiple imputation (MI) (Larsen, 2011; von Hippel, 2016). A practical benefit of FIML compared to MI is the stability of estimation across uses, whereas multiple imputation depends on stochastic sampling and will yield a (slightly) different estimate every time.

Moreover, combining information across different imputations can be challenging, although this has been automated for SEM with lavaan via the auxiliary package ‘semTools’ (Jorgensen et al., 2015).

Once model estimation has finished (which may take anywhere from fractions of a second to days), a wide range of model fit indices are available to assess model fit (Kline, 2011; Schermelleh-Engel et al., 2003). Generally, these metrics quantify the deviation between the observed and implied covariance matrix. Model fit metrics include a simple test of deviation from perfect model fit (the chi square test), indices that compare the degree to which the proposed model better fits the data (e.g. the CFI and TLI) compared to some baseline model (which typically is a model in which there are no correlations between measurements; and good fit represents the degree to which covariation in the empirical data is reliably modelled), and measures that quantify some standardized measures of the deviation between the observed and implied covariance matrices (e.g. SRMR or the RMSEA). Fit indices can be affected by a range of model and data properties including sample size, measurement quality, estimation method, misspecification and more (Fan, Thompson, & Wang, 1999; McNeish, An, & Hancock, 2017; Moshagen & Erdfelder, 2016). Competing models can be compared using traditional likelihood ratio test if models are nested (Neale, 2000), or specialized version of the LRT for non-nested models (Merkle, You, & Preacher, 2016). Other approaches to model comparison include the use of model fit indices such as CFI and RMSEA (Usami et al., 2016), or information based metrics such as the AIC and BIC (Aho, Derryberry, & Peterson, 2014; Wagenmakers & Farrell, 2004). A relatively new question inspired by cognitive neuroscience will be how to best conduct model selection and model comparison in procedures such as voxelwise modelling from brain image data (Madhyastha et al., n.d.) which may require hundreds or thousands of model comparisons or extended measurement models of spatially correlated observations – This at present remains an open challenge.

Debates concerning the optimal way to assess and interpret model fit, which thresholds to use or how to best compare models are wide ranging and beyond the scope of this tutorial, for further details see (Barrett, 2007; Fan, Thompson, & Wang, 1999; Hayduk, Cummings, Boadu, Pazderka-Robinson, & Boulianne, 2007; Schermelleh-Engel, Moosbrugger, & Müller, 2003; Steiger, 2007). Common advice includes reporting multiple (types of) fit indices to allow for a more holistic assessment such as reporting raw chi^2, CFI and RMSEA. (Schermelleh-Engel et al., 2003). Recommended sources for a wide range of (longitudinal) SEM topics include McArdle, (2009), Newsom (2015), Hoyle (2014), Little, (2013), (Voelkle & Oud, 2015), Driver et al. (2016) and Voelkle (2007), as well as the tutorials cited above. Other useful resources are SEM-oriented email groups such as SEMNET (http://www2.gsu.edu/~mkteer/semnet.html) or package focused help groups (e.g. https://groups.google.com/forum/#!forum/lavaan)

### 4.1.2 Convergence and improper solutions

Although SEM in general and LCS in particular cover a broad and flexible range of models and techniques, these techniques have various limitations. Below we outline a subset of commonly faced challenges. After specifying an LCS model, researchers will use a particular estimation procedure, typically Maximum Likelihood, to provide estimates for the parameters in the model. However, in contrast to simpler methods such as t-tests and simple regressions, model estimation may fail to converge. Common causes of a failure to converge include small sample sizes, overly complex models, poor starting values, inappropriate estimators, large amounts of missing data, data input errors, or misspecified models (Fan et al., 1999; Jackson, 2007; Wothke, 1993). A particular challenge in the context of latent change score models (and closely related models such as linear mixed models) is that of estimating or constraining variances terms (close) to 0 (for instance, constraining the variance to 0 in the simplest univariate latent change score model will generally lead to non-convergence even if the change scores between T1 and T2 are identical across individuals). Classical estimation methods such as Maximum Likelihood have relatively high non-convergence rates in such scenarios, and non-convergence has often (erroneously) been interpreted as evidence that the model is necessarily inappropriate. Moreover, the likelihood ratio test may not behave appropriately in scenarios including such ‘boundary values’ (Stoel et al., 2006). One promising solution is Bayesian estimation, which has been suggested to have considerably less estimation problems (Eager & Roy, 2017; Merkle & Rosseel, 2015; B. Muthén & Asparouhov, 2012; Van Erp, Mulder, & Oberski, 2017). Secondly, even if estimation does converge, model fit may be ‘improper’ in various ways. Such improper solutions may include negative variances, standardized estimates that (far) exceed 1 (sometimes referred to as ‘Heywood cases’, but see Jöreskog, 1999), or matrices that are ‘non-positive definite’ (Wothke, 1993). No unique cause underlies these problems, nor does a single solution exist that applies in all cases, but various remedies may help. These including providing plausible starting values for parameters to aid estimation, increasing sample sizes, using a different estimator, or constraining parameters to 0 or to (in)equality where appropriate - for example, variances which are estimated just below zero might be constrained to zero so that they remain within appropriate bounds. For further reading on challenges and solutions of model convergence and inappropriate solutions we recommend (Eager & Roy, 2017; Fan et al., 1999; Gerbing & Anderson, 1987; Wothke, 1993)

### 4.1.3 Power and sample size

A challenge closely related to model fit and model comparison is that of statistical power and the associated question of sample size (Cohen, 1988). One often encounters rules of thumb (such as “one should have 20 subjects per SEM parameter”), which typically are misleading and never capture the full story. Here, we advise against such heuristics. When designing a longitudinal study, there are many more design decisions that directly affect statistical power, such as indicator reliability, true effect size, or the number and spacing of measurement occasions (Brandmaier, von Oertzen, Ghisletta, & Lindenberger, 2015), and strategies exist to improve power without necessarily increasing sample size (Hansen & Collins, 1994). Longitudinal models have successfully been fit to as few as 22 subjects (Curran, Obeidat, & Losardo, 2010), but as for all statistical approaches, larger sample sizes will generally lead to more robust inferences. Although factors determining statistical power in latent growth models are reasonably well understood (Hertzog, Lindenberger, Ghisletta, & Oertzen, 2006; Hertzog, von Oertzen, Ghisletta, & Lindenberger, 2008; Oertzen, Hertzog, Lindenberger, & Ghisletta, 2010; Rast & Hofer, 2014; von Oertzen & Brandmaier, 2013; von Oertzen, Brandmaier, & Tsang, 2015), we know of no empirical simulation studies that may serve as guidelines for the power of (bivariate) LCSM. Currently, statistical power for a specific LCSM design (including a hypothesized true effect and sample size) can be approximated mathematically (Satorra & Saris, 1985) or by computer-intensive simulation (L. K. Muthén & Muthén, 2002). For a more general approach to the question of model selection, model comparison and parameter recovery we have included a flexible, general script that can allow users to generate a dataset under known conditions from a given model, and fit one or more models to this dataset. This simple approach should allow anyone to examine compare parameter recovery, convergence rates, statistical power, model selection and model fit under a range of effect sizes, sample sizes and missingness to facilitate appropriate study planning.

### 4.1.4 Inference and causality

Once model convergence and adequate model fit are obtained, the most daunting step is that of (causal) inference: What may and may not be concluded based on the results? One goal of SEM is to test predictions derived from causal hypotheses about the process that generated the data, represented as a model. That is, although SEM (nor any other correlation-based technique) cannot directly demonstrate causality or causal processes, it can be used as a statistical tool for deriving model-based predictions of causal hypotheses, and examine the extent to which the data disconfirms these hypotheses (Bollen & Diamantopoulos, 2015; Pearl, 2000). However, inferring causality is, unsurprisingly, non-trivial. A first and most fundamental challenge, not specific to SEM, is that of *model equivalence,* known in philosophy of science as the ‘the underdetermination of theory by data’ (e.g. Newton-Smith & Lukes, 1978). In the context of SEM, it has been shown formally that any observed data pattern is compatible with many different data generating mechanisms (Raykov & Penev, 1999). In other words, even if a model fits well in a sufficiently large dataset, that in and of itself is not conclusive evidence for the (causal) hypotheses posited by the model. Moreover, in a longitudinal context, modelling choices and omitted variables can affect, and even spuriously invert, causal direction and temporal ordering (Usami et al., 2016) as well as the magnitude (Voelkle & Oud, 2015) of effects. Although recent years have seen a consistent trend away from causal language (with recommendations to move away from the term ‘causal modelling’ as shorthand for SEM (Kline, 2011), for a spirited defence as well as a historically informed overview of the causal history and foundations of SEM, see Pearl (2012). The most commonly accepted solution is that model inferences, including causality, should come from a convergence of robust empirical evidence guided by theoretical motivations, and ideally be validated by interventions whenever possible.

Finally, although SEM is commonly used as a technique to test whether data is (provisionally) compatible with a particular (causal) hypothesis, in practice SEM spans a range of approaches from almost entirely exploratory to confirmatory. In the context of LCS models, non-trivial misfit may be amended by model re-specification to achieve acceptable fit. One approach to improve model fit is the examination of ‘modification indices’ – The expected improvement in model fit if a currently constrained parameter was freely estimated. As this is generally purely data-driven, this practice may adversely affect interpretability and generalization to independent datasets, so should be exercised with caution (MacCallum, Roznowski, & Necowitz, 1992). Other approaches include the addition of cross-loadings, the elimination of non-significant structural paths, constraining or equating parameters or the exclusion of poorly performing measurement indicators. All of those changes may be defensible, but researchers should be explicit about any modifications that were made purely to improve model fit, so as to be able to assess the evidence in favour of the ‘final’ model appropriately (Bentler, 2007; MacCallum et al., 1992).

### 4.2 LCS versus alternative models

Structural equation modelling is an extremely general framework to study differences and change, and shares foundations with other analytic approaches. Previous work has shown similarities and even equivalences with other analytical traditions such as multilevel- and linear mixed modelling (Bauer, 2003; Curran, 2003; Rovine & Molenaar, 2001). However, even when models are mathematically equivalent in principle, they may still diverge in practical terms, such as ease of model specification and common defaults - McNeish & Matta (2017) examine in which situations linear mixed models versus SEM approaches are the likely to be the better choice. An overarching treatment by Voelkle (Voelkle, 2007) has shown how a wide variety of analytical techniques ranging from *t*-tests to MANOVAs can be (re)written as special cases of the latent growth curve model (which is itself a special case of the latent change score model). For instance, (Coman et al., 2013) shows how a basic LCS is a special case of the paired *t*-test. Similarly, simple forms of the bivariate latent change score model can be rewritten as a special case of a cross-lagged panel model, namely the recently proposed random-intercept cross-lagged panel model (Hamaker et al., 2015), and the autoregressive cross-lagged factor model is equivalent to a latent change score model when slope factor scores are equivalent across individuals (Usami et al., 2016).

Grimm (2007) used three popular longitudinal models, the bivariate latent growth curve model, the latent growth curve with a time-varying covariate, and the bivariate dual change score growth model, to examine the same dataset concerning the relation between depression and academic achievement. Although the three models yielded different results, Grimm (2007) illustrates how each of the three approaches answer slightly different developmental questions, illustrating the importance of McArdle’s question posed at the beginning of this article: ‘When thinking about any repeated measures analysis it is best to ask first, what is your model for change?’ (McArdle, 2009, p. 579). Alternative longitudinal SEM approaches that address specific questions with differing strengths and weaknesses include the autoregressive latent trajectory (ALT) model (Bollen & Curran, 2004), survival models (Newsom, 2015, chapter 12), continuous time models (Driver et al., 2016), simplex models (Newsom, 2015), the incorporation of definition variables (Mehta & West, 2000), regime switching LCS models (Chow, Grimm, Filteau, Dolan, & McArdle, 2013), latent trait-state models (Steyer, Schmitt, & Eid, 1999), and extensions of latent curve models including structured residuals and time-varying covariates (Curran, Howard, Bainter, Lane, & McGinley, 2014). For a general introduction to longitudinal SEM approaches we recommend the recent book by Newsom (2015).

Direct comparisons of LCS models to competing models exist but are relatively rare. Usami et al. (2016) compared the LCS to the auto-regressive cross-lagged factor (ARCL) model, and showed lower levels of bias in the parameter estimates of the LCS model, depending on the number of time points and sample size, but slightly more power for the ARCL model (due to decreased standard errors). Notably, they observed that model selection was best when using the less conventional approach of comparison model fit indices (RMSEA and CFI) as opposed to likelihood ratio tests or information indices. Using simulation studies, McArdle & Hamagami, (2001a) compared the bivariate dual change score model to a Multilevel Change Score Regression Model under a range of known data generating processes. They showed that the multilevel regression model performed adequately only under a range of restrictive conditions including no missing data, an absence of error terms and no bivariate coupling. The bivariate dual change score model on the other hand was able to accurately recover parameter estimates under a range of missingness conditions, even up to the extreme case of cross-sectional data (i.e. one timepoint per individual), as long as data was Missing Completely at Random (p. 233), illustrating the robustness and flexibility of LCS models. Compared to simpler, more traditional techniques LCS and related models more natural accommodate a range of commonly encountered research challenges, including missing data, unequal spacing, time-varying covariates, and latent and manifest group comparisons which may aid in the nature, direction and precision of statistical inferences in studying dynamic processes (Curran et al., 2010). Many developmental hypotheses can be cast as a special case of the LCS, but researchers should always carefully considered whether a given model best captures their central developmental hypotheses compared to other analytical approaches.

## 5 Fitting Latent Change Score models using open source software

A wide array of tools exist to fit longitudinal SEM models, ranging from modules within popular statistical tools (e.g., AMOS within SPSS; Arbuckle, 2010) to dedicated SEM software (e.g., Mplus; Muthén & Muthén, 2005). We focus on two freely available tools: The package lavaan (Rosseel, 2012) within R and a standalone, GUI-based tool Ωnyx (von Oertzen et al., 2015).

### 5.1 Lavaan

R (R Development Core Team, 2016) is a powerful programming language with a rapidly growing user community dedicated to data analysis and visualisation. Several excellent interactive introductions to R exist, including http://tryr.codeschool.com/ or http://swirlstats.com/. The core strength of R is the wide range of packages dedicated to addressing specific challenges (more than 10,000 as of February 2017), implementing statistical techniques, visualisation and more. Several packages dedicated to SEM exist, including OpenMx (Boker et al., 2011) which allows for a high degree of model specification flexibility, but relatively complex syntax), the sem package (Fox, 2006), Bayesian SEM (blavaan, Merkle & Rosseel, 2015), regularized SEM for complex models (regsem, Jacobucci, Grimm, & McArdle, 2016) and even a new package dedicated specifically to specific subtypes of longitudinal SEM (RAMpath, Zhang et al., 2015).

We will focus on lavaan (Rosseel, 2012) as this is a highly popular and versatile tool for modelling various structural equation models, including longitudinal models (Rosseel, 2013). Lavaan syntax consists of multiple lines specifying relations among variables using different operators for e.g. factor loadings (‘=~’), regressions (‘~’), (co)variances (‘~~’), and means or intercepts (‘~1’). In the syntax below (Figure 6) we specify a simple, univariate latent change score model, estimating five key parameters (in bold).

**Figure 6:**
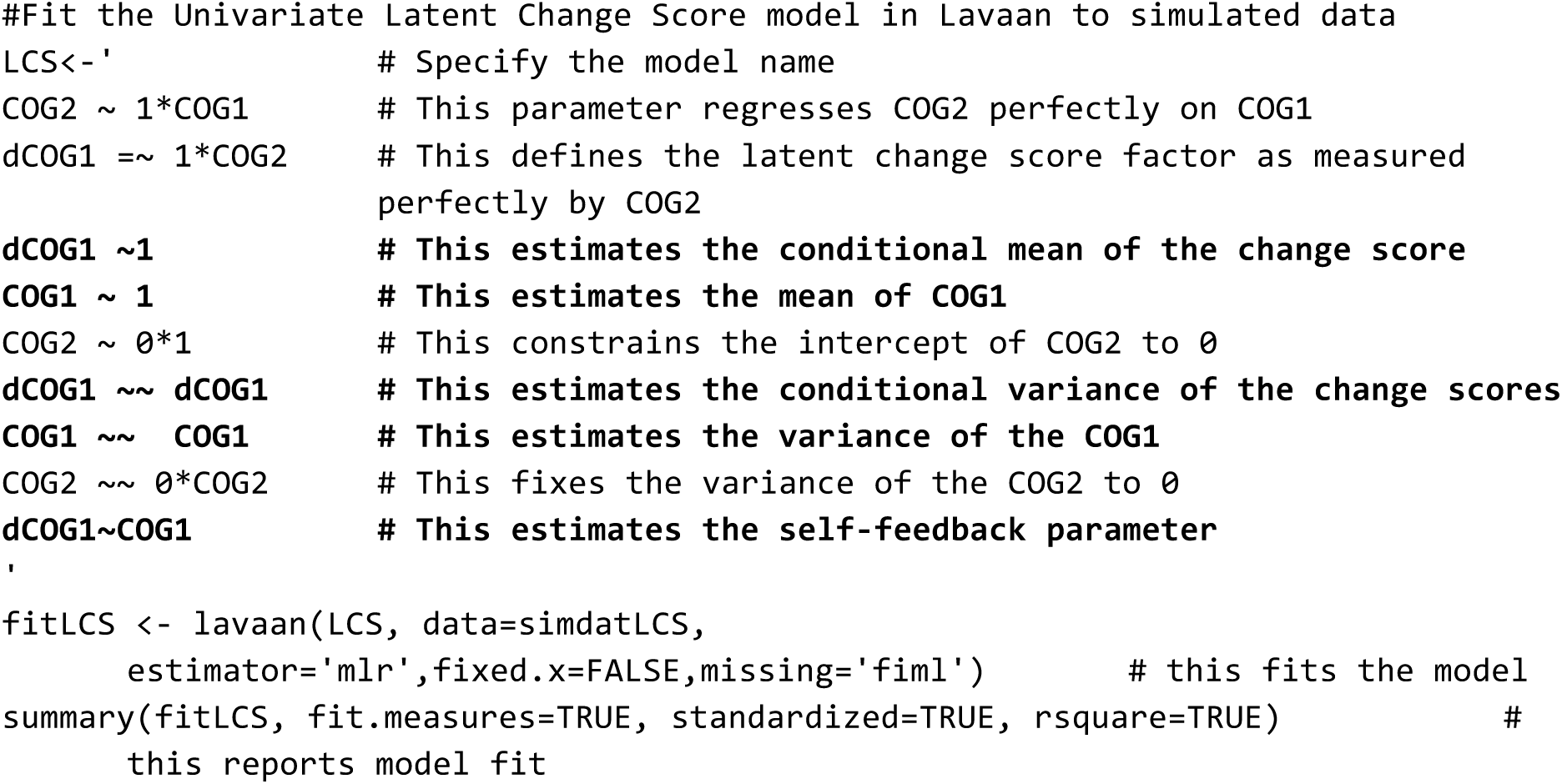
Lavaan example syntax. Comments in R are preceded by #. Key LCS parameters are boldfaced.

The lavaan syntax and simulated data for all five model types discussed above is available online^https://osf.io/4bpmq/?view_only=5b07ead0ef5147b4af2261cb864eca32^. These scripts install and load the relevant packages if needed, simulate data according to given set of parameter values, visualize raw data and fit the model. For a simulated data object called ‘simdatLCS’, the syntax above fits a simple Univariate Latent Change Score model with a Yuan-Bentler correction for non-normality (‘estimator=’mlr’), and full information maximum likelihood to deal with missing data (‘missing=’fiml’). In Appendix A we provide a step-by-step instruction to fit models using R or Ωnyx (see below).

### 5.2 Ωnyx

Although syntax-centred methods for SEM are most common, new users may prefer a more visual, path model based approach (e.g. AMOS, Arbuckle, 2010). One powerful tool is Ωnyx (von Oertzen et al., 2015), a freely available graphical modelling software for creating and estimating various structural equation models. At the time of writing, we used the most recent public version (Ωnyx 1.0-937), available from http://onyx.brandmaier.de/. Ωnyx provides a purely graphical modelling environment without a model syntax level, that is, models are simply drawn as path diagrams. As soon as datasets are loaded within a model, estimation starts on-the-fly and parameter estimates will be directly shown in the model diagram. In addition to its easy-to-use interface, a particular strength of Ωnyx is its capability of generating model syntax for other programs, such as Lavaan (Rosseel, 2012) OpenMx (Boker et al., 2011), or Mplus (L. K. Muthén & Muthén, 2005). The focus on the graphical interface makes Ωnyx especially useful for beginners who want to get a basic comprehension of SEM, but also for more advanced users who either want to transition to other SEM programs or need to produce diagrams for presentations or manuscripts. Finally, Ωnyx provides template models for commonly used models, reducing time needed to set up standard models to a minimum. Here, we will give a brief introduction on how the Ωnyx graphical user interface works (also see and, specifically, how LCSMs can be set up and estimated.

The idea behind Ωnyx is a little different to typical editors. The main menu is virtually empty (with the exception of basic load and save functions) and there is neither a tool ribbon (e.g., as in Microsoft Word) or a toolbox (e.g., as in Adobe Photoshop) to access functions. Instead, Ωnyx relies heavily on context-menus that are accessible with right mouse-clicks. A double-click performs a context-specific default action. For example, when Ωnyx is started, the empty Ωnyx desktop is shown. A right-click on the desktop opens a new model frame, which is a container for a SEM. Alternatively, a double-click on the desktop creates a model frame (see Figure 7 for an example of the interface). In Appendix A, we provide a step by step guide to fitting an existing model to data within Ωnyx, as well as a step-by-step explanation how to specify a new model from scratch.5

**Figure 7:**
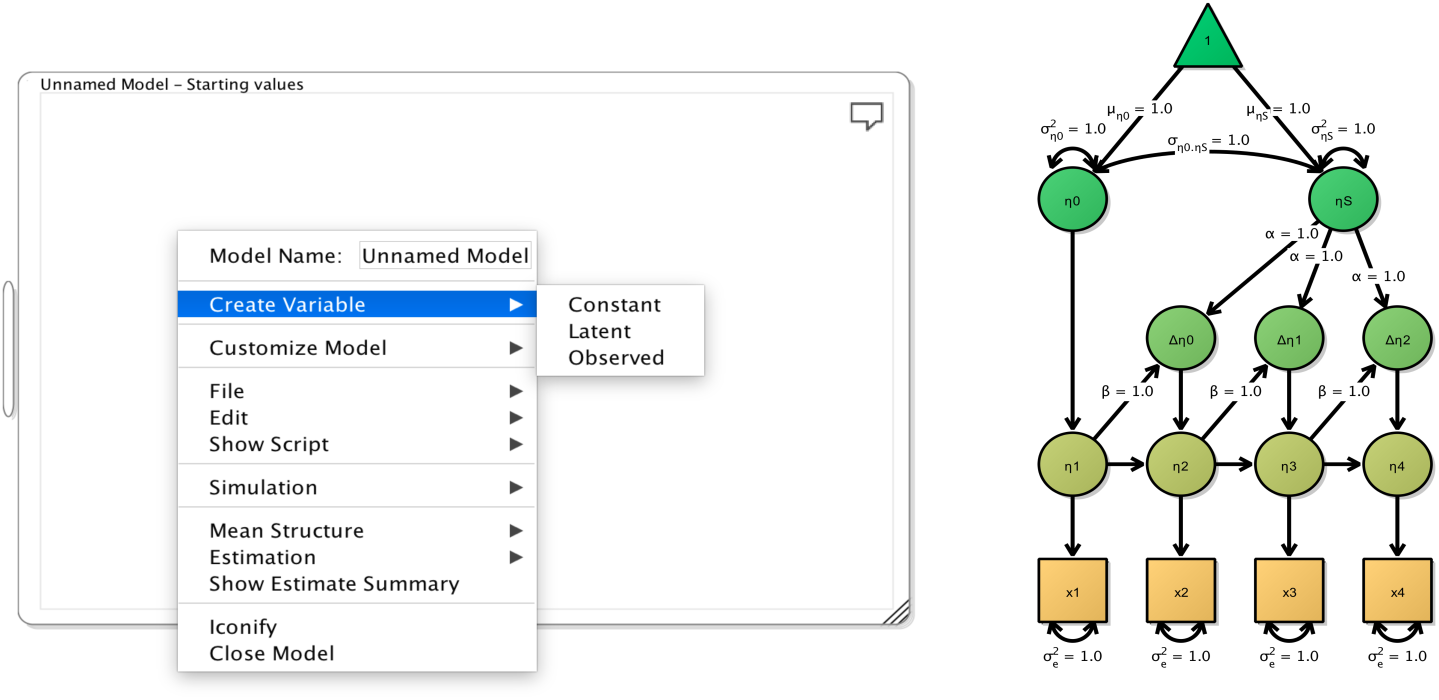
Ωnyx interface

### 5.3 Developing intuitions about change using an interactive Shiny app

Above we explained the basics of LCS models, including graphical representations. Although these examples are relatively easy to understand, one challenge with complex dynamic models is that it can be hard to intuit what the consequences of changes in various parameters might be. To ameliorate this problem, we have built an interactive online tool using the R package Shiny (Chang, Cheng, & Allaire, 2016). This tool allows researchers to modify the key parameters of interest for three key models (univariate latent change score, bivariate latent change score, and bivariate dual change score) in an interactive fashion and examine the consequences for the observed scores. Figure 8 illustrates our shiny interface, which can be found at http://brandmaier.de/shiny/sample-apps/SimLCS_app/^3^. The drop-down menu at the top can be used to select one of three models, and the sliders can be used to tweak individual parameters. Changing the key parameters causes the underlying simulation to be modified on the fly, and the panels at the bottom visualize the raw data simulated.

**Figure 8:**
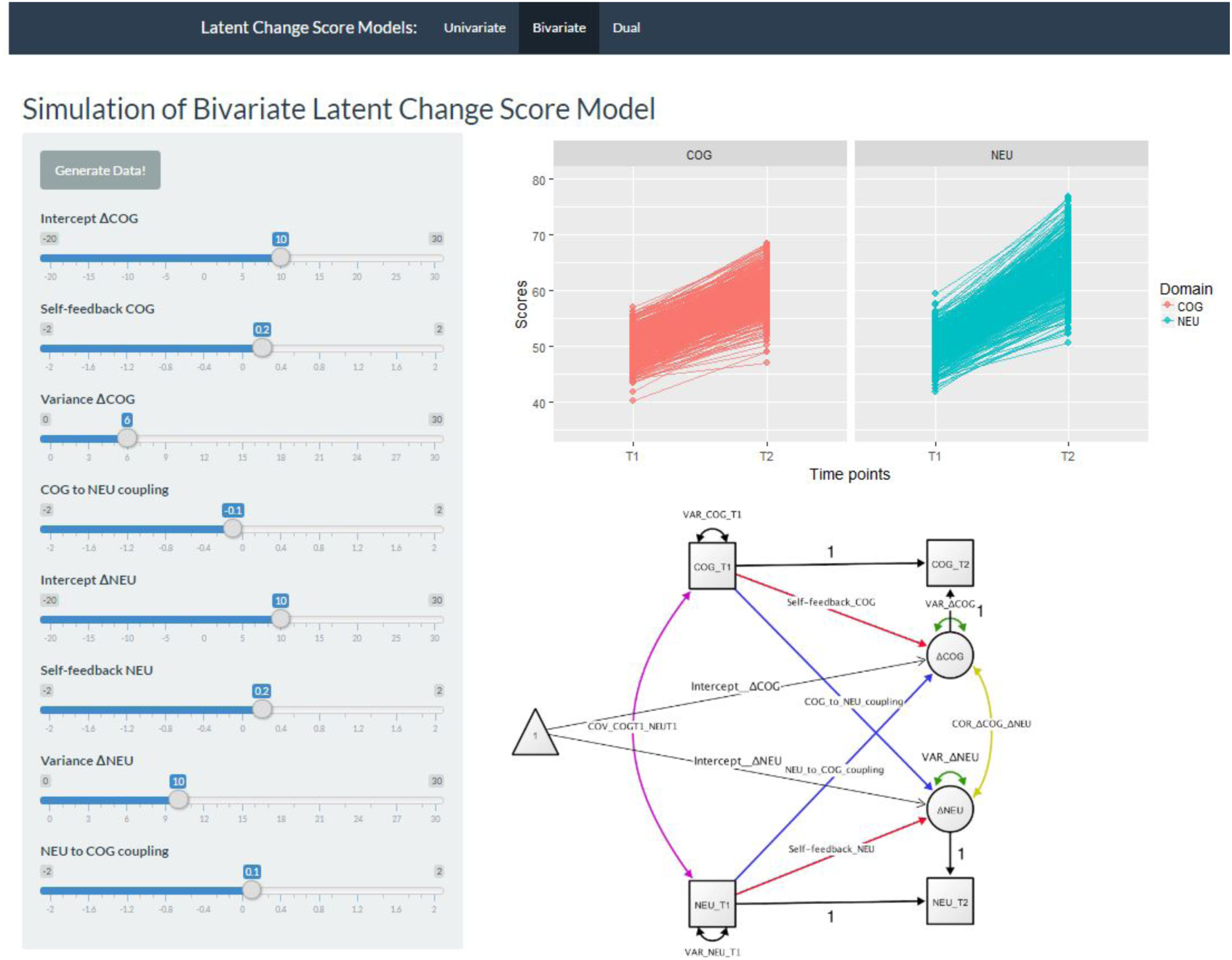
Shiny interface. At the top users can select from three different latent change score models (Univariate, bivariate or dual). At the left, users can modify key parameters and select ‘generate data’ to simulate data with a given parametrisation. On the right, the raw data as well as the path model will shown. This allows users to form an intuition of the effect of dynamic coupling. For instance, it can illustrate that even in the absence of significant changes within a domain, a coupling parameter from another domain can cause significant increases or decreases

The underlying code can easily be accessed and modified, such that researchers can tailor our code to their specific research design. Our hope is that this tool will prove useful in developing intuitions about dynamic co-occurring processes of development and change.

## 6 Examples

Below we illustrate the flexibility of Latent Change Score modelling by describing two empirical examples. First, we describe cognitive (processing speed) and neural (white matter fractional anisotropy) changes from a training intervention study in younger and older adults. Second, we describe group differences in structural changes (i.e. cortical thinning) in a developmental study of (late) adolescents aged 14-24. These applications illustrate the types of questions naturally accommodated by latent change score models.

### 6.1 Correlated change in high intensity training intervention: The COGITO sample

The first illustration comes from data on the COGITO project (Schmiedek et al., 2014), a high-intensity (100 day) training intervention with pre and post-tests cognitive scores for 204 adults: 101 young (age M=25.11, SD=2.7, range=20-31) and 103 old (age M=70.78, SD=4.15, range=64-80).

We examine changes between pre- and post-test scores on a latent variable of processing speed, measured by three standardized tests from the Berlin Intelligence Structure test measured on two occasions (see Schmiedek et al. (2010) for more details). The neural measure of interest is fractional anisotropy in the sensory subregion of the Corpus Callosum (see Lövdén et al., 2010 for more details - note in our exploratory analysis this subregion gave the most stable results and so it was used for our illustration). Longitudinal neuroimaging data was available for a subset of 32 people (20 younger, 12 older adults). We fit the model to the entire sample using Full Information Maximum Likelihood estimation, maximizing the use of our sample and decreasing bias compared to complete case analysis. However, the neural parameters should be interpreted with a level of caution commensurate to the modest sample size. See Lövdén et al. (2014) for further discussion regarding the benefits of FIML in such a context and Enders (2001) for more general discussion of FIML. First, we test a multiple indicator univariate latent change score model (the same type of model as shown in Figure 3). This univariate (only processing speed) multiple indicator (a latent variable of processing speed is specified) latent change score model fits the data well: *χ*^2^(12)= 15.052, *P*= 0.239, RMSEA= 0.035 [0.000 0.084], CFI=0.996, SRMR=0.028, Yuan-Bentler scaling factor= 0.969). Inspection of key parameters shows that scores increased between pre- and post-test (the intercept of the change factor= 0.224, se=0.031, Z=7.24), there were significant individual differences in gains (variance parameter for the latent change score: est=0.120, se=0.019, Z=6.5), but the rate of improvement did not depend on the starting point: est=-0.069, se=0.054, Z=-1.32). Next, we include a neural measure, namely Fractional Anisotropy in the sensory region of the Corpus Callosum measured pre- and post-test, to fit a *bivariate* (neural and behaviour) *multiple indicator* (we include a measurement model) *latent change score model* shown in Figure 9.

**Figure 9:**
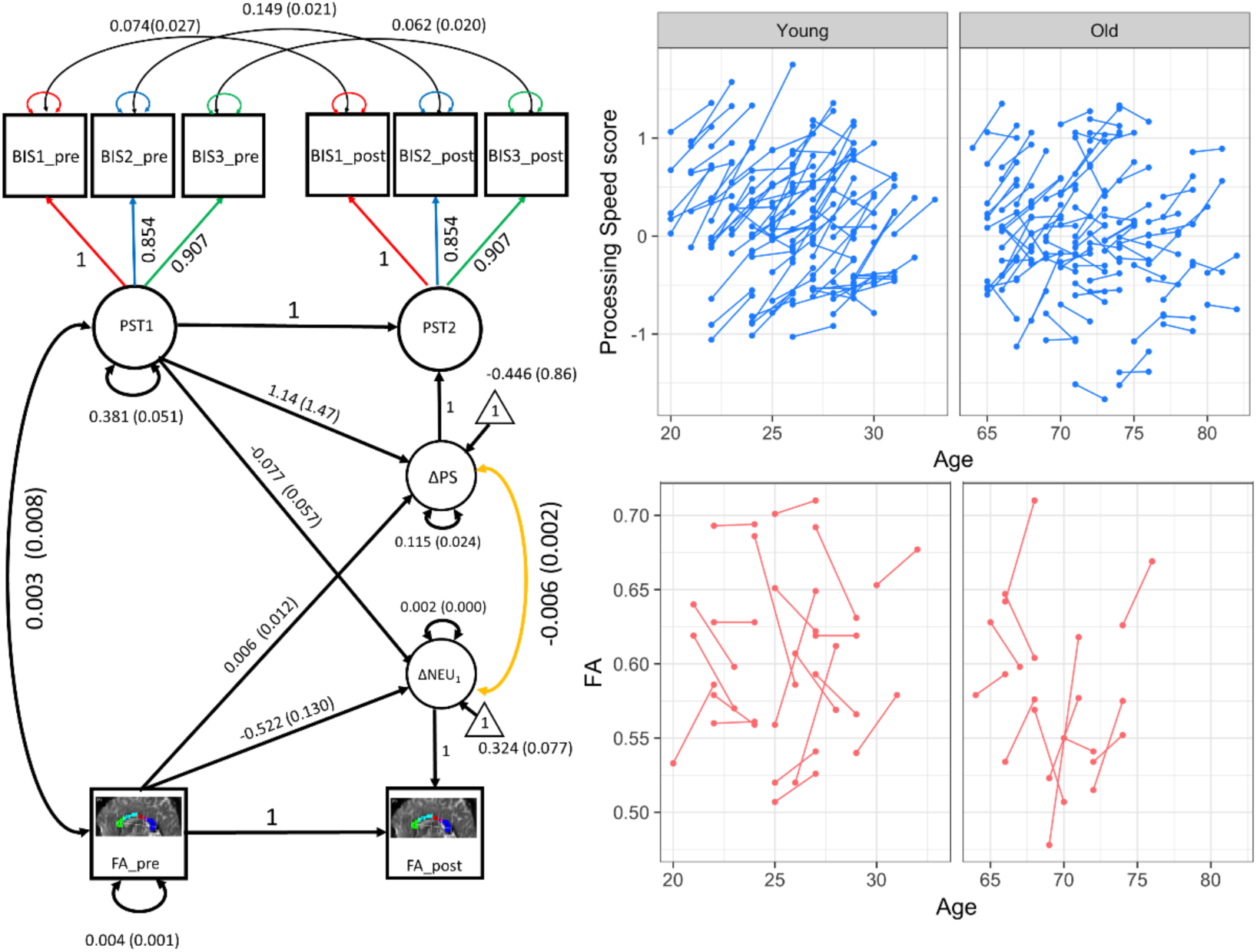
COGITO correlated change in processing speed and white matter plasticity. The panels on the left show the fitted model, parameter estimates and standard errors. The latent variable Processing Speed is measured by three subtests of the Berlin Intelligence Structure test (BIS1-BIS3) measured before (pre) and after (post) an intensive training intervention (see Schmiedek et al., 2010). Observed variable means and variance estimates are omitted for visual clarity. The panels on the right show the raw scores changing across two occasions. The raw scores are plotted on separate panels to accommodate the age gap, but the model is estimated for the population as a whole.

We next test the evidence for four possible brain-behaviour relationships: *Covariance* (are scores on processing speed at T1 *correlated with* white matter structure at T1?), neural measures as leading variable (do differences in white matter integrity at T1 affect the rate of cognitive training gains?), cognition as leading variable (do processing speed scores at T1 predict degree of white matter plasticity between T1 and T2?) and/or *correlated change* (is the degree of improvement in the cognitive domain correlated with the degree of white matter change in individuals?). Figure 9 shows the full model, as well as the changes in processing speed factor scores (top right) and fractional anisotropy (bottom right) (note we artificially expanded the interval between testing intervals for visual clarity). First, we find that model fit is good: *χ*^2^(20)= 24.773, *P*= 0.21, RMSEA= 0.034 [90% CI: 0.000 0.072], CFI=0.995, SRMR=0.083, Yuan-Bentler scaling factor= 1.021. The full model is shown in Figure 9. Inspection of the four parameters of interest, reflecting the four possible brain-behaviour relationships outlined above, shows evidence (only) for *correlated change*. In other words, those with greater gains in processing speed were, on average, those with *less* positive change in fractional anisotropy after taking into account the other dynamic parameters (est=-0.006, se=0.002, z=-2.631). Although counterintuitive, a similar pattern was also observed in Bender, Prindle, Brandmaier, & Raz (2015) who observed negative correlation between age-related declines in episodic memory and white matter integrity, such that a greater decrease in fractional anisotropy was associated with greater improvement in episodic memory (whereas at T1 FA and episodic memory were positively correlated). This illustration shows how LCS models can be used to simultaneously estimate four rather distinct brain-behaviour relationships over time.

### 6.1 Multigroup analysis of prefrontal structural change in late adolescence: The NSPN cohort

As the example in the Cogito sample shows, LCS offers a simple and powerful way to test distinct dynamic pathways within a single LCS model. However, investigations in Developmental Cognitive Neuroscience are often concerned with differences between groups (e.g. gender, treatment vs. controls, psychopathology vs. healthy controls, low vs high SES etc.). Such questions are best addressed by means of *multigroup modelling*. Here we illustrate a multigroup model to compare structural changes in a group of adolescents. Data for this is drawn from the Neuroscience in Psychiatry Network (NSPN), a cohort that studies development in adolescents (see also Kiddle et al., 2017; Kievit et al., 2017; Whitaker et al., 2016) Here we illustrate a multigroup model to compare structural brain change in a group of adolescents. Previous work suggests differences in the temporal development of the frontal cortex, with boys generally maturing later than girls (Giedd et al., 2012; Ziegler, Ridgway, Blakemore, Ashburner, & Penny, 2017), possibly as a consequence of differences in sensitivity to hormone levels (Bramen et al., 2012).

For our analysis, we focus on volumetric changes in the frontal pole. This region is part of the frontal lobe, which is often discussed with respect to the speed of maturational changes and its purported role in controlling higher cognitive functions and risk taking behaviour (e.g. Crone & Dahl, 2012; Johnson, 2011; Mills et al., 2014).

Our sample consisted of 176 individuals, mean age = 18.84, range 14.3-24.9, 82 girls, scanned on two occasions (average interval: M=1.24 years, SD= 0.33 years). We fit a multiple indicator univariate latent change score model, with volume of the frontal pole (FP) using the neuromorphometrics atlas as the key variable (see Figure 10D for an illustration). Our measurement model consisted of volumetric estimates of the left and right FP measured on two occasions (for more details on the structural processing pipeline, see Appendix B). We can use the framework of multigroup models to investigate whether there is evidence for differences between the two groups in the key parameters of interest. The four parameters of interest are the mean of the frontal factor (reflecting mean volumes at T1), the intercept of the change factor (reflecting the rate of change), the variance of the latent change scores (reflecting individual differences in rates of FP change) and the covariance between T1 and rate of change. To do so, we employ the method of equality constrained Likelihood Ratio tests, by comparing a model where some parameter of interest is constrained to be the same across the two groups with a model where the parameter is allowed to be free. The difference in model fit under the null hypothesis is chi-square distributed with a df equivalent to the difference in numbers of parameters being constrained. In other words, if a parameter of interest is the same (or highly similar) between two groups, the chi-square test will fail to be rejected, suggesting the more parsimonious model is sufficient.

**Figure 10:**
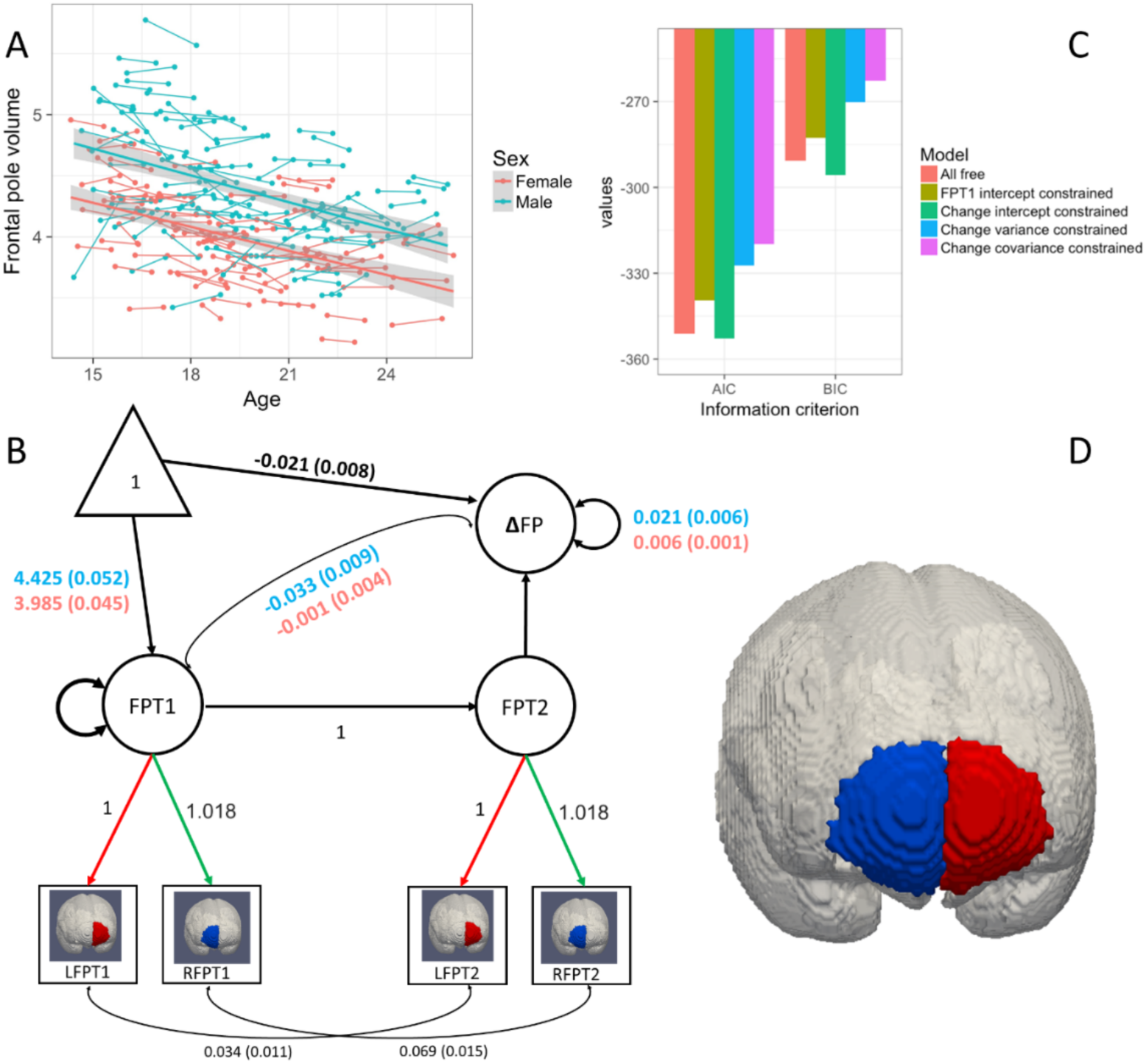
NSPN: Differential variability in frontal lobe thinning. Panel A shows longitudinal development in frontal structure. Panel B shows the model fit for the best model. Where parameters are different between groups we show male estimates in blue, female in red. Panel C shows the AIC and BIC of the free versus constrained models – in all cases only one parameter is constrained to equality and compared to the ‘all free’ model. Panel D shows the left and right frontal poles of the neuromorphometrics atlas used in our analysis. See Appendix B for more details on the imaging pipeline.

First, we fit a model where all measurement model parameters (constraining all factor loadings and residual (co)variances) are constrained to be equal across males and females, but all other parameters are free to vary between the sexes. This model fit the data well: *χ*^2^(9)= 8.929, *P*= 0.44, RMSEA= 0.00 [0.000 0.120], CFI=1, SRMR=0.021, Yuan-Bentler scaling factor= .983). Next, we explored which (if any) of the four parameters above differed between the sexes. If a parameter is different between the groups, constraining it to be equal should result in a significant decrease in model fit. Using the likelihood ratio test, we observe significant decreases in model fit by constraining the mean of frontal lobe volume at T1 to be equal across the sexes (χ2Δ = 38.01, dfΔ =2, p=<0.0001). Inspection of parameter estimates shows, unsurprisingly, greater FP volume in males, compatible with either larger brains, delayed cortical thinning, or a combination of the two. Contrary to expectations, constraining the intercept of the change scores did not lead to a significant decrease in fit (χ2Δ = 0.31889, dfΔ =2, p=0.57), indicating an absence of reliable differences in cortical thinning. However, constraining the *variance* of change scores to be equal did result in a significant drop in fit (χ2Δ = 49.319, dfΔ =2, p=<0.0001), with males showing greater individual differences in rates of thinning than females. Finally, constraining the covariance between frontal volume and change scores also led to a drop in model fit, with males showing a stronger (negative) association between volume at T1 and rate of change (compatible with the hypotheses of delayed development in males).

Together, this suggests that there are considerable differences in frontal development between males and females in (late) adolescence: Although males and females show similar rates of cortical thinning, males show greater initial volume, greater individual differences in thinning and a stronger association between initial volume and rate of thinning. Notably, the parameters where the evidence for sex differences is strongest (e.g. variance and covariance in change scores) are not parameters often studied using conventional techniques such as paired t-tests (other than as a statistical assumption such as equality of variances). Conversely, the parameter that would be the key focus with traditional techniques (i.e. group differences in change scores) does not show differences. Figure 10 shows the temporal development of FP structure between sexes, information based model comparison and parameter estimates for the full model (with different estimates for males and females where required).

## 7 Conclusion

In this tutorial, we introduce the powerful framework of Latent Change Score modelling whose deployment can be an invaluable aid for developmental cognitive neuroscience. It is our hope that more widespread employment of these powerful techniques will aid the developmental cognitive neuroscientific community. Adopting the statistical techniques we outline in tandem with the more widespread availability of large, longitudinal, cohorts of developing adolescents will allow researchers to more fully address questions of interest, as well as inspire new questions and approaches. The approach we outline puts renewed emphasis on the value of longitudinal over cross-sectional data in addressing developmental questions.

## Appendix A

Creating and fitting Latent Change Score models using R and Ωnyx

In the associated folder you will find code (file extensions. R), data (file extensions. csv) and Ωnyx model files (file extensions. xml) for five different latent change score models. Below we outline how to specify and fit models using lavaan and R.

### 1 Analyse data using R and lavaan

-Install R (https://cran.r-project.org/)

-Install Rstudio (recommended) (https://www.rstudio.com/)

-Open the relevant lavaan file (e.g. ‘1_ULCS.R’)

-Install the appropriate packages by uncommenting (e.g. lines 17-19 in 1_ULCS.R)

-Select and run lines 30-60 to create a simulated dataset with given parameters

-Select and run lines 85-92 to visualize the raw data

-Select and run lines 65-80 to fit the latent change score model

-Run line 81 to examine model fit and parameters

-If so desired, modify the parameters in lines 47-53 to examine the consequences for the raw data and model fit

### 2 Fit a model to existing dataset using Ωnyx

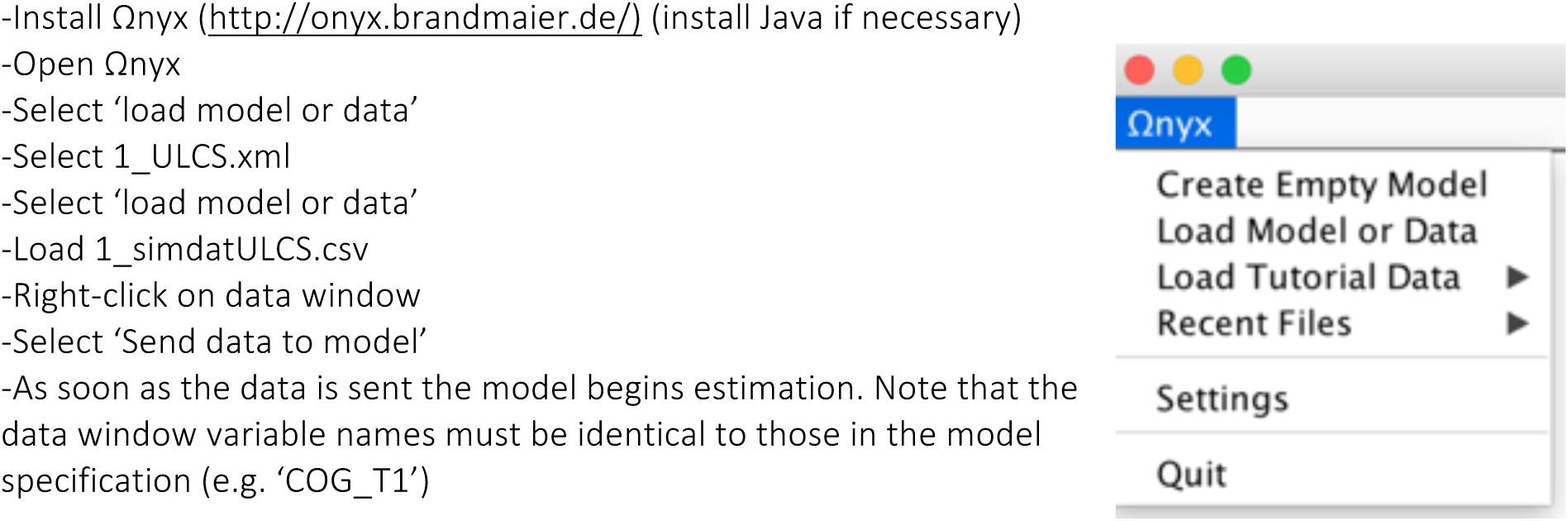

### 3 Creating a new model using Ωnyx

To create variables in the model, right-click on the model frame and choose ‘add variable’ to add an observed, latent or constant variable. Alternatively, users can double-click on the model frame to create an observed variable. Existing variables can be moved by left-dragging (press left mouse button and move mouse while button pressed). Double-clicking while holding down the SHIFT button creates a latent variable. Regression paths (single headed) are drawn by right drags, that is, by pressing the right mouse button on a variable and releasing the button only when the mouse was moved to a second variable. Covariance paths (double headed) are drawn by holding SHIFT while releasing a path. Variance paths are created by creating a covariance path from a variable to itself. By default, path values are fixed to one. Paths values can be changed either by right-clicking a path and entering a new value in the context menu or by moving the mouse over a path and directly typing the desired value. Path can be defined to represent a freely estimated parameter by right-clicking a path and choosing “Free Parameter”. Using the context menu, parameters can be renamed and starting values can be given. Observed variables can be either associated with a data column or not. This is indicated by observed variables either having a grey box (not linked) or a black box (linked data). To link a variable with data, one can load a dataset from an existing file and simply drag the variables onto the Ωnyx representation.

As an alternative to the manual creation of a change score model, Ωnyx provides a wizard for quick model specification, even for more advanced models. Users can right-click on the Ωnyx desktop and choose, among other models, ‘Create new LGCM’ (for a linear growth curve model) or ‘Create new DCSM’ (for a dual change score model), which can then be modified as desired. The LCSM wizard allows you to specify the number of time points. Once the model is specified, users can simply drag and drop data (e.g. columns in a .csv file) from an existing data file onto the appropriate observed variables in the model by selecting ‘load data’ from the main drop-down menu. As soon as variables are added the program starts estimation.

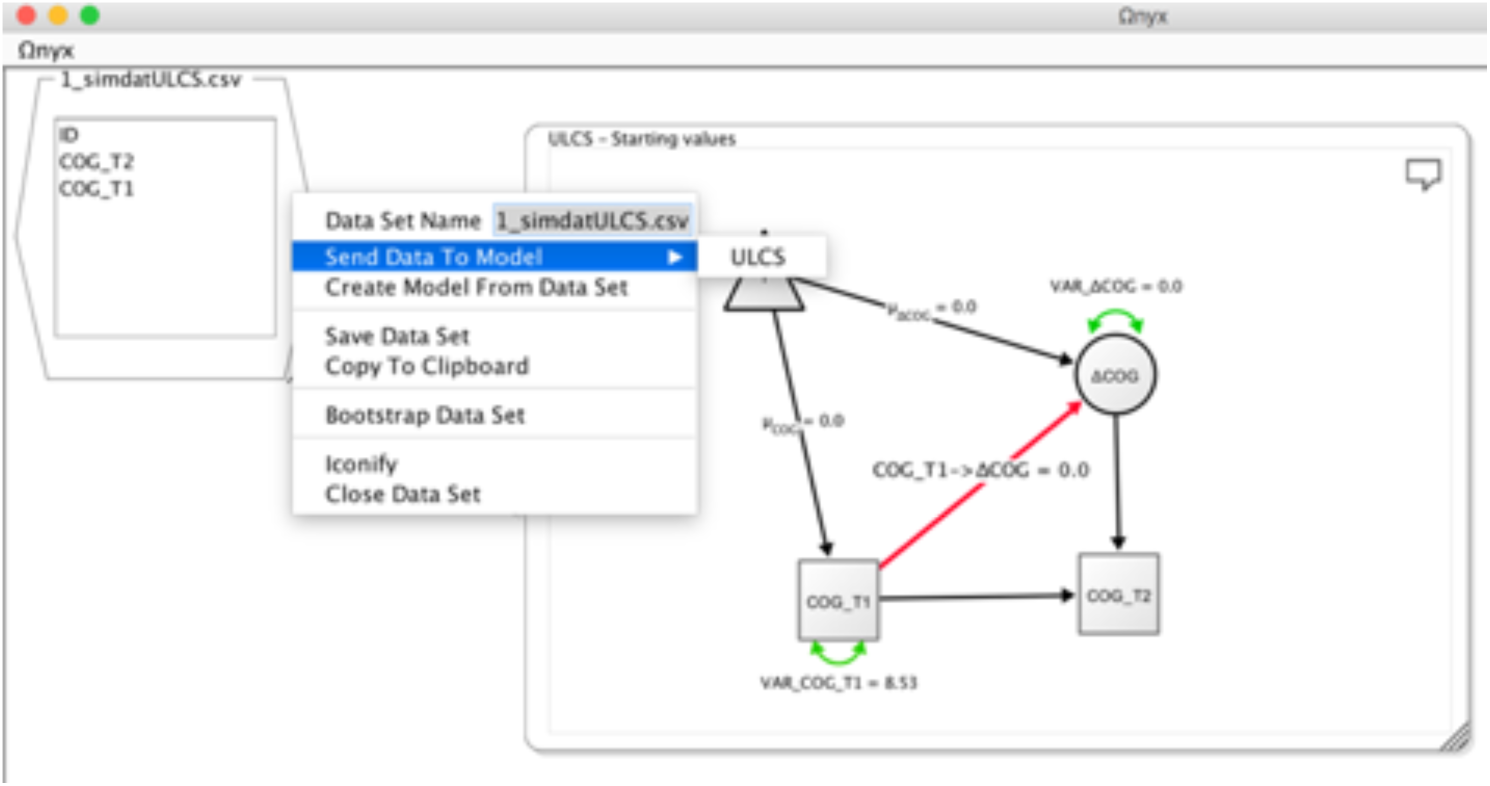

### Advanced features of Ωnyx

As noted before, Ωnyx provides various functions to export models to other SEM programs. For example, if users wish to use more complicated modelling approaches (e.g., ordinal outcomes, non-linear constraints), Ωnyx can quickly create a graphical model specification which then can be exported to another SEM program that allows greater flexibility in modelling. Ωnyx also exports diagrams as bitmap graphics (JPG, PNG) or vector graphics (PDF, EPS) and even creates LaTeX-representations of diagrams based on the tikz package. Ωnyx comes with different visual style templates that can be applied to a model with a single click. To choose among styles, right-click on a model frame and choose Customize Model → Apply Diagram Style, or hit CTRL+L to cycle through styles in the active model frame. Ωnyx may be used to simulate data from the model-implied means and covariance matrix. To this end, choose “Simulation->Start Simulation” in the context menu of a model frame. Ωnyx allows parameter names to be defined in a pseudo LaTeX input style, which allows users to use greek symbols (e.g., \alpha, \beta, \gamma,…), subscripts (e.g., \epsilon_i) and superscripts (e.g., \sigma^2).

## Appendix B

### Structural Imaging and Map Generation

Brain scans were acquired using the MPM protocol (Weiskopf et al., 2013) on three 3T whole-body MRI systems (Magnetom TIM Trio; VB17 software version; Siemens Healthcare) located in Cambridge and London. Between-site reliability of MRI procedures was assessed in a pilot study scanning five healthy volunteers at each site. The between-site bias was found to be less than 3% and the between-site coefficient of variation was less than 8% for both longitudinal relaxation rate (R1) and MT parameters (Weiskopf et al., 2013) Isotropic 1 mm MT images were quantified in Matlab (2014b, The MathWorks, Inc.) using SPM12 r6685 (Wellcome Trust Centre for Neuroimaging, London, UK, http://www.fil.ion.ucl.ac.uk/spm), the Voxel-Based Quantification (VBQ) toolbox for SPM (Callaghan et al., 2014) and custom made tools.

### Longitudinal Image Processing and Feature Extraction

All further image processing steps of MT maps were performed in SPM12. Longitudinal morphometry was used to assess macroscopic brain maturational changes. Since longitudinal imaging is prone to artefacts due to registration inconsistency, scanner inconsistencies and age-related deformations of the brains, it requires sophisticated processing pipelines in order to detect the changes of interest and achieve unbiased results.

First, we applied symmetric diffeomorphic registration for longitudinal MRI (Ashburner & Ridgway, 2012) combining non-linear diffeomorphic and rigid-body registration and correction for intensity inhomogeneity artefacts. The optimization is realized within one integrated generative model and provides consistent estimates of within-subject brain deformations over the study period. The registration model also creates a midpoint image for each subject and the corresponding deformation fields for every individual scan.

Second, we applied SPM12's Computational Anatomy Toolbox (CAT, r955, Structural Imaging Group, http://dbm.neuro.uni-jena.de/cat12/) segmentation to each subject's midpoint image, which assumes every voxel to be drawn from an unknown mixture of gray matter (GM), white matter (WM), cerebrospinal fluid (CSF). Earlier results showed that MT maps are highly suitable for automated segmentation in multi-subject morphometric studies, showing improved GM tissue contrast in subcortical structures (Helms, Draganski, Frackowiak, Ashburner, & Weiskopf, 2009). This applied segmentation procedure contains partial volume estimation (PVE) to account for mixed voxels with two tissue types (Tohka, Zijdenbos, & Evans, 2004). The CAT algorithm is based on an adaptive maximum a posteriori (AMAP) approach (Rajapakse, Giedd, & Rapoport, 1997) and subsequent application of a hidden Markov random field model (Cuadra, Cammoun, Butz, Cuisenaire, & Thiran, 2005). Importantly, the applied AMAP estimation does not rely on tissue priors, which overcomes potential bias due to the application of inappropriate tissue priors in young maturing subjects with different to adult brain anatomy.

Third, nonlinear template generation and image registration was performed on the individual midpoint GM and WM tissue maps using DARTEL registration and the template was registered to MNI space using an affine transform (Ashburner, 2007). Consecutively longitudinal normalised tissue segments from all subjects and time-points were modulated by Jacobian determinants accounting for local tissue volume differences across subjects and within-subject changes over time. In order to detect stronger deviations due to potential segmentation or normalization errors, we included a quality check using covariance-based sample inhomogeneity measures implemented in the CAT toolbox to exclude subjects with extremal values and/or severe artefacts.

Fourth, neuromorphometrics atlas was used to assess gray matter density in the bilateral frontal poles after Gaussian smoothing with 6 mm full width at half maximum. The atlas was based on maximum probability tissue labels derived in the MICCAI 2012 Grand Challenge and Workshop on Multi-Atlas Labeling with data originating from the OASIS project (http://www.oasis-brains.org/) and the atlas provided by Neuromorphometrics, Inc. (http://Neuromorphometrics.com/) under academic subscription.

All longitudinal features for subsequent structural equation modelling were obtained using the above steps. Since common processing pipelines of longitudinal and cross-sectional data is different, this can introduce biases (Bernal-Rusiel et al., 2013). We initially focussed on the largest fully longitudinally processed subsample of the NSPN dataset available, with 202 subjects having at least 2 scans per person. After rigorous quality control of all processed imaging data, scans from 26 subjects had to be discarded due to artefacts, resulting in a finally analyzed brain features from 373 scans from 176 subjects (2.12 scans/person). Although a small subset of individuals had a third intermediate scan, for modelling purposes here we use only the first and last scan.

## Acknowledgements

We would like to thank Delia Fuhrmann, Janne Adolf, Rasmus Berggren and Stuart Ritchie for comments on a previous draft of this manuscript, as well as readers of our preprint who reached out with valuable suggestions for improvement.

## Principal investigators

Edward Bullmore (CI from 01/01/2017) Raymond Dolan

Ian Goodyer (CI until 01/01/2017) Peter Fonagy

Peter Jones

## NSPN (funded) staff

Michael Moutoussis

Tobias Hauser

Petra Vértes

Kirstie Whitaker

Gita Prabhu

Laura Villis

Junaid Bhatti

Becky Inkster

Cinly Ooi

Barry Widmer

Ayesha Alrumaithi

Sarah Birt

Kalia Cleridou

Hina Dadabhoy

Sian Granville

Elizabeth Harding

Alexandra Hopkins

Daniel Isaacs

Janchai King

Danae Kokorikou

Harriet Mills

Ciara O’Donnell

Sara Pantaleone

## Affiliated scientists

Pasco Fearon

Anne-Laura van Harmelen

Rogier Kievit

## Funding sources

RAK is supported by the Sir Henry Wellcome Trust (grant number 107392/Z/15/Z) and the UK Medical Research Council Programme (MC-A060-5PR61). The NSPN cohort was supported by a strategic award by the Wellcome Trust to the University of Cambridge and University College London (095844/Z/11/Z).

**Conflict of interest statement:**

E.T.B. is employed half-time by the University of Cambridge and half-time by GlaxoSmithKline; he holds stock in GlaxoSmithKline.

## Footnotes

1 It is worth noting that such hypotheses of temporal precedence in measurable properties do not imply a dualist perspective on mental and physical processes (cf. Kievit et al., 2011). They do suggest scientifically relevant distinctions can be made with implications for interpretation, the likely consequences of interventions and early detection of non-typical development.

2 Although it should be noted that SEM software packages can vary in their default model specifications, so users should always be aware of these modelling assumptions

3 Note: some firewalls block the app. A zipped folder that contains all scripts and can be run locally is available on http://brandmaier.de/shiny/sample-apps/SimLCS_app/

## References

Ahmed, S. P., Bittencourt-Hewitt, A., & Sebastian, C. L. (2015). Neurocognitive bases of emotion regulation development in adolescence. Developmental Cognitive Neuroscience, 15, 11–25. http://doi.org/10.1016/j.dcn.2015.07.006

Aho, K., Derryberry, D., & Peterson, T. (2014). Model selection for ecologists: the worldviews of AIC and BIC. Ecology, 95(3), 631–636. http://doi.org/10.1890/13-1452.1

Arbuckle, J. L. (2010). IBM SPSS ^®^ Amos ^TM^ 19 User's Guide. Retrieved from http://amosdevelopment.com

Ashburner, J. (2007). A fast diffeomorphic image registration algorithm. NeuroImage, 38(1), 95–113. http://doi.org/10.1016/j.neuroimage.2007.07.007

Ashburner, J., & Ridgway, G. R. (2012). Symmetric diffeomorphic modeling of longitudinal structural MRI. Frontiers in Neuroscience, 6, 197. http://doi.org/10.3389/fnins.2012.00197

Baltes, P., Reese, H. W., & Nesselroade, J. R. (1977). Life-Span Developmental Psychology: Introduction to Research Methods. Monterey: Brooks-Cole. Retrieved from https://www.questia.com/library/3760900/life-span-developmental-psychology-introduction-to

Baraldi, A. N., & Enders, C. K. (2010). An introduction to modern missing data analyses. Journal of School Psychology, 48, 5–37. http://doi.org/10.1016/j.jsp.2009.10.001

Barrett, P. (2007). Structural equation modelling: Adjudging model fit. Personality and Individual Differences, 42(5), 815–824. http://doi.org/10.1016/j.paid.2006.09.018

Bauer, D. J. (2003). Estimating Multilevel Linear Models as Structural Equation Models. Journal of Educational and Behavioral Statistics, 28(2), 135–167. http://doi.org/10.3102/10769986028002135

Bauer, D. J. (2007). Observations on the Use of Growth Mixture Models in Psychological Research. Multivariate Behavioral Research, 42(4), 757–786. http://doi.org/10.1080/00273170701710338

Bender, A. R., Prindle, J. J., Brandmaier, A. M., & Raz, N. (2015). White matter and memory in healthy adults: Coupled changes over two years. NeuroImage. http://doi.org/10.1016/j.neuroimage.2015.10.085

Bengtsson, S. L., Nagy, Z., Skare, S., Forsman, L., Forssberg, H., & Ullén, F. (2005). Extensive piano practicing has regionally specific effects on white matter development. Nature Neuroscience, 8(9), 1148–50. http://doi.org/10.1038/nn1516

Bentler, P. M. (2007). On tests and indices for evaluating structural models. Personality and Individual Differences, 42(5), 825–829. http://doi.org/10.1016/j.paid.2006.09.024

Bernal-Rusiel, J. L., Greve, D. N., Reuter, M., Fischl, B., Sabuncu, M. R., & Alzheimer's Disease Neuroimaging Initiative. (2013). Statistical analysis of longitudinal neuroimage data with Linear Mixed Effects models. NeuroImage, 66, 249–260. http://doi.org/10.1016/j.neuroimage.2012.10.065

Boker, S., Neale, M., Maes, H., Wilde, M., Spiegel, M., Brick, T., … Fox, J. (2011). OpenMx: An Open Source Extended Structural Equation Modeling Framework. Psychometrika, 76(2), 306–317. http://doi.org/10.1007/s11336-010-9200-6

Bollen, K. A. (1989). Structural Equations with Latent variables. New York: Wiley.

Bollen, K. A., & Curran, P. J. (2004). Autoregressive Latent Trajectory (ALT) Models A Synthesis of Two Traditions. Sociological Methods & Research, 32(3), 336–383. http://doi.org/10.1177/0049124103260222

Bollen, K. A., & Diamantopoulos, A. (2015). In Defense of Causal-Formative Indicators: A Minority Report. Psychological Methods. http://doi.org/10.1037/met0000056

Bramen, J. E., Hranilovich, J. A., Dahl, R. E., Chen, J., Rosso, C., Forbes, E. E., … Dinov, I. (2012). Sex Matters during Adolescence: Testosterone-Related Cortical Thickness Maturation Differs between Boys and Girls. PLoS ONE, 7(3), e33850. http://doi.org/10.1371/journal.pone.0033850

Brandmaier, A. M., Prindle, J. J., McArdle, J. J., & Lindenberger, U. (2016). Theory-guided exploration with structural equation model forests. Psychological Methods, 21(4), 566–582. http://doi.org/10.1037/met0000090

Brandmaier, A. M., von Oertzen, T., McArdle, J. J., & Lindenberger, U. (2013). Structural equation model trees. Psychological Methods, 18(1), 71–86. http://doi.org/10.1037/a0030001

Callaghan, M. F., Freund, P., Draganski, B., Anderson, E., Cappelletti, M., Chowdhury, R., … Weiskopf, N. (2014). Widespread age-related differences in the human brain microstructure revealed by quantitative magnetic resonance imaging. Neurobiology of Aging, 35(8), 1862–72. http://doi.org/10.1016/j.neurobiolaging.2014.02.008

Chang, W., Cheng, J., & Allaire, J. J. (2016). shiny: Web Application Framework for R.

Cheung, G. W., & Rensvold, R. B. (2002). Evaluating Goodness-of-Fit Indexes for Testing Measurement Invariance. Structural Equation Modeling: A Multidisciplinary Journal, 9(2), 233–255. http://doi.org/10.1207/S15328007SEM0902_5

Chow, S.-M., Grimm, K. J., Filteau, G., Dolan, C. V., & McArdle, J. J. (2013). Regime-Switching Bivariate Dual Change Score Model. Multivariate Behavioral Research, 48(4), 463–502. http://doi.org/10.1080/00273171.2013.787870

Coman, E. N., Picho, K., McArdle, J. J., Villagra, V., Dierker, L., & Iordache, E. (2013). The paired t-test as a simple latent change score model. Frontiers in Psychology, 4, 738. http://doi.org/10.3389/fpsyg.2013.00738

Crone, E. A., & Dahl, R. E. (2012). Understanding adolescence as a period of social-affective engagement and goal flexibility. Nature Reviews Neuroscience, 13(9), 636–650. http://doi.org/10.1038/nrn3313

Cuadra, M. B., Cammoun, L., Butz, T., Cuisenaire, O., & Thiran, J.-P. (2005). Comparison and validation of tissue modelization and statistical classification methods in T1-weighted MR brain images. IEEE Transactions on Medical Imaging, 24(12), 1548–65. http://doi.org/10.1109/TMI.2005.857652

Curran, P. J. (2003). Have Multilevel Models Been Structural Equation Models All Along? Multivariate Behavioral Research, 38(4), 529–569. http://doi.org/10.1207/s15327906mbr3804_5

Curran, P. J., Howard, A. L., Bainter, S. a., Lane, S. T., & McGinley, J. S. (2014). The separation of between-person and within-person components of individual change over time: A latent curve model with structured residuals. Journal of Consulting and Clinical Psychology, 82(5), 879–894. http://doi.org/10.1037/a0035297

Curran, P. J., Obeidat, K., & Losardo, D. (2010). Twelve Frequently Asked Questions About Growth Curve Modeling. Journal of Cognition and Development: Official Journal of the Cognitive Development Society, 11(2), 121–136. http://doi.org/10.1080/15248371003699969

Driver, C. C., Oud, J. H. L., & Voelkle, M. C. (2016). Continuous Time Structural Equation Modelling With R Package ctsem. Journal of Statistical Software.

Eager, C., & Roy, J. (2017). Mixed Effects Models are Sometimes Terrible. Retrieved from http://arxiv.org/abs/1701.04858

Enders, C. K. (2001). A Primer on Maximum Likelihood Algorithms Available for Use With Missing Data. Structural Equation Modeling: A Multidisciplinary Journal, 8(1), 128–141. http://doi.org/10.1207/S15328007SEM0801_7

Fan, X., Thompson, B., & Wang, L. (1999). Effects of sample size, estimation methods, and model specification on structural equation modeling fit indexes. Structural Equation Modeling: A Multidisciplinary Journal, 6(1), 56–83. http://doi.org/10.1080/10705519909540119

Ferrer, E., Balluerka, N., & Widaman, K. F. (2008). Factorial Invariance and the Specification of Second-Order Latent Growth Models. Methodology, 4(1), 22–36. http://doi.org/10.1027/1614-2241.4.1.22

Ferrer, E., & McArdle, J. J. (2004). An Experimental Analysis of Dynamic Hypotheses About Cognitive Abilities and Achievement From Childhood to Early Adulthood. Developmental Psychology, 40(6), 935–952.

Ferrer, E., Shaywitz, B. A., Holahan, J. M., Marchione, K., & Shaywitz, S. E. (2010). Uncoupling of reading and IQ over time: empirical evidence for a definition of dyslexia. Psychological Science, 21(1), 93–101. http://doi.org/10.1177/0956797609354084

Fox, J. (2006). TEACHER'S CORNER: Structural Equation Modeling With the sem Package in R. Structural Equation Modeling: A Multidisciplinary Journal, 13(3), 465–486. http://doi.org/10.1207/s15328007sem1303_7

Gerbing, D. W., & Anderson, J. C. (1987). Improper solutions in the analysis of covariance structures: Their interpretability and a comparison of alternate respecifications. Psychometrika, 52(1), 99–111. http://doi.org/10.1007/BF02293958

Gesell, A. (1929). Maturation and infant behavior pattern. Psychological Review, 36(4), 307–319. http://doi.org/10.1037/h0075379

Ghisletta, P., & Lindenberger, U. (2003). Age-Based Structural Dynamics Between Perceptual Speed and Knowledge in the Berlin Aging Study: Direct Evidence for Ability Dedifferentiation in Old Age. Psychology and Aging, 18(4), 696–713. http://doi.org/10.1037/0882-7974.18.4.696

Ghisletta, P., & McArdle, J. J. (2012). Teacher's Corner: Latent Curve Models and Latent Change Score Models Estimated in R. Structural Equation Modeling: A Multidisciplinary Journal, 19(4), 651–682. http://doi.org/10.1080/10705511.2012.713275

Giedd, J. N., Raznahan, A., Mills, K. L., Lenroot, R. K., Cohen, S. B., Lombardo, M., … Hare, T. (2012). Review: magnetic resonance imaging of male/female differences in human adolescent brain anatomy. Biology of Sex Differences, 3(1), 19. http://doi.org/10.1186/2042-6410-3-19

Gorbach, T., Pudas, S., Lundquist, A., Orädd, G., Josefsson, M., Salami, A., … Nyberg, L. (2016). Longitudinal association between hippocampus atrophy and episodic-memory decline. Neurobiology of Aging. http://doi.org/10.1016/j.neurobiolaging.2016.12.002

Grimm, K. J. (2007). Multivariate longitudinal methods for studying developmental relationships between depression and academic achievement. International Journal of Behavioral Development, 31(4), 328–339. http://doi.org/10.1177/0165025407077754

Hamaker, E. L., Kuiper, R. M., & Grasman, R. P. P. P. (2015). A critique of the cross-lagged panel model. Psychological Methods, 20(1), 102–116. http://doi.org/10.1037/a0038889

Hansen, W. B., & Collins, L. M. (1994). Seven ways to increase power without increasing N. NIDA Research Monograph, 142, 184–95. Retrieved from http://www.ncbi.nlm.nih.gov/pubmed/9243537

Hayduk, L., Cummings, G., Boadu, K., Pazderka-Robinson, H., & Boulianne, S. (2007). Testing! testing! one, two, three - Testing the theory in structural equation models! Personality and Individual Differences, 42(5), 841–850. http://doi.org/10.1016/j.paid.2006.10.001

Helms, G., Draganski, B., Frackowiak, R., Ashburner, J., & Weiskopf, N. (2009). Improved segmentation of deep brain grey matter structures using magnetization transfer (MT) parameter maps. NeuroImage, 47(1), 194–8. http://doi.org/10.1016/j.neuroimage.2009.03.053

Hertzog, C., Lindenberger, U., Ghisletta, P., & Oertzen, T. von. (2006). On the power of multivariate latent growth curve models to detect correlated change. Psychological Methods, 11(3), 244–252. Retrieved from http://psycnet.apa.orgjournals/met/11/3/244

Hertzog, C., von Oertzen, T., Ghisletta, P., & Lindenberger, U. (2008). Evaluating the Power of Latent Growth Curve Models to Detect Individual Differences in Change. Structural Equation Modeling: A Multidisciplinary Journal, 15(4), 541–563. http://doi.org/10.1080/10705510802338983

Hoyle, R. H. (2014). Handbook of structural equation modeling. (R. H. Hoyle, Ed.). Guilford press.

Jackson, D. L. (2007). The Effect of the Number of Observations per Parameter in Misspecified Confirmatory Factor Analytic Models. Structural Equation Modeling: A Multidisciplinary Journal, 14(1), 48–76. http://doi.org/10.1080/10705510709336736

Jacobucci, R., Grimm, K. J., & McArdle, J. J. (2016). Regularized Structural Equation Modeling. Structural Equation Modeling: A Multidisciplinary Journal, 23(4), 555–566. http://doi.org/10.1080/10705511.2016.1154793

Jajodia, & Archana. (2012). Dynamic structural equation models of change. Routledge/Taylor & Francis Group.

Johnson, M. H. (2011). Interactive Specialization: A domain-general framework for human functional brain development? Developmental Cognitive Neuroscience, 1(1), 7–21. http://doi.org/10.1016/j.dcn.2010.07.003

Jöreskog, K. G. (1999). How Large Can a Standardized Coefficient be? Retrieved from http://www.ssicentral.com/lisrel/techdocs/HowLargeCanaStandardizedCoefficientbe.pdf

Jorgensen, T. D., Pornprasertmanit, S., Miller, P., Schoemann, A., Rosseel, Y., Quick, C., … Al, E. (2015). Package “semTools”.

Kiddle, B., Inkster, B., Prabhu, G., Moutoussis, M., Whitaker, K., Consortium, N., … Jones, P. (2017). The NSPN 2400 Cohort: a developmental sample supporting the Wellcome Trust Neuroscience in Psychiatry Network. http://doi.org/10.17863/CAM.11026

Kievit, R. A., Davis, S. W., Mitchell, D., Taylor, J. R., Duncan, J., Cam-CAN, & Henson, R. N. (2014). Distinct aspects of frontal lobe structure mediate age-related differences in fluid intelligence and multitasking. Nature Communications, 5(5658), 1–10.

Kievit, R. A., Frankenhuis, W. E., Waldorp, L. J., & Borsboom, D. (2013). Simpson's paradox in psychological science: a practical guide. Frontiers in Psychology, 4, 513. http://doi.org/10.3389/fpsyg.2013.00513

Kievit, R. A., Lindenberger, U., Goodyer, I. M., Jones, P. B., Fonagy, P., Bullmore, E. T., … Dolan, R. J. (2017). Mutualistic coupling between vocabulary and reasoning supports cognitive development during late adolescence and early adulthood. Psychological Science.

Kievit, R. A., Romeijn, J.-W., Waldorp, L. J., Wicherts, J. M., Scholte, H. S., & Borsboom, D. (2011). Mind the Gap: A Psychometric Approach to the Reduction Problem. Psychological Inquiry, 22(2), 67–87. http://doi.org/10.1080/1047840X.2011.550181

Kline, R. B. (2011). Principles and Practice of Structural Equation Modeling. Retrieved from http://books.google.com/books?hl=nl&lr=&id=mGf3Ex59AX0C&pgis=1

Larsen, R. (2011). Missing Data Imputation versus Full Information Maximum Likelihood with Second-Level Dependencies. Structural Equation Modeling: A Multidisciplinary Journal, 18(4), 649–662. http://doi.org/10.1080/10705511.2011.607721

Lindenberger, U., & Poetter, U. (1998). The Complex Nature of Unique and Shared Effects in Hierarchical Linear Regression: Implications for Developmental Psychology, 3(2), 218–230.

Lindenberger, U., von Oertzen, T., Ghisletta, P., & Hertzog, C. (2011). Cross-sectional age variance extraction: What's change got to do with it? Psychology and Aging, 26(1), 34–47. Retrieved from http://psycnet.apa.orgjournals/pag/26/1/34

Little, T. D. (2013). Longitudinal structural equation modeling.

Little, T. D., Lindenberger, U., & Nesselroade, J. R. (1999). On selecting indicators for multivariate measurement and modeling with latent variables: When “good” indicators are bad and “bad” indicators are good. Psychological Methods, 4(2), 192–211. http://doi.org/10.1037/1082-989X.4.2.192

Lövdén, M., Bodammer, N. C., Kühn, S., Kaufmann, J., Schütze, H., Tempelmann, C., … Lindenberger, U. (2010). Experience-dependent plasticity of white-matter microstructure extends into old age. Neuropsychologia, 48(13), 3878–3883. http://doi.org/10.1016/j.neuropsychologia.2010.08.026

Lövdén, M., Köhncke, Y., Laukka, E. J., Kalpouzos, G., Salami, A., Li, T.-Q., … Bäckman, L. (2014). Changes in perceptual speed and white matter microstructure in the corticospinal tract are associated in very old age. NeuroImage, 102P2, 520–530. http://doi.org/10.1016/j.neuroimage.2014.08.020

Lövdén, M., Li, S.-C., Shing, Y. L., & Lindenberger, U. (2007). Within-person trial-to-trial variability precedes and predicts cognitive decline in old and very old age: longitudinal data from the Berlin Aging Study. Neuropsychologia, 45(12), 2827–38. http://doi.org/10.1016/j.neuropsychologia.2007.05.005

Maass, A., Düzel, S., Goerke, M., Becke, A., Sobieray, U., Neumann, K., … Düzel, E. (2015). Vascular hippocampal plasticity after aerobic exercise in older adults. Molecular Psychiatry, 20(5), 585–593. http://doi.org/10.1038/mp.2014.114

MacCallum, R. C., Roznowski, M., & Necowitz, L. B. (1992). Model modifications in covariance structure analysis: The problem of capitalization on chance. Psychological Bulletin, 111(3), 490–504. http://doi.org/10.1037/0033-2909.111.3.490

Madhyastha, T., Peverill, M., Koh, N., McCabe, C., Flournoy, J., Mills, K. L., … McLaughlin, K. (n.d.). Current methods and limitations for longitudinal fMRI analysis across development. Developmental Cognitive Neuroscience.

Malone, P. S., Lansford, J. E., Castellino, D. R., Berlin, L. J., Dodge, K. A., Bates, J. E., & Pettit, G. S. (2004). Divorce and Child Behavior Problems: Applying Latent Change Score Models to Life Event Data. Structural Equation Modeling: A Multidisciplinary Journal, 11(3), 401–423. http://doi.org/10.1207/s15328007sem1103_6

McArdle, J. J. (1994). Structural Factor Analysis Experiments with Incomplete Data. Multivariate Behavioral Research, 29(4), 409–454. http://doi.org/10.1207/s15327906mbr2904_5

McArdle, J. J. (2008). Latent Variable Modeling of Differences and Changes with Longitudinal Data. Retrieved from http://www.annualreviews.org/doi/abs/10.1146/annurev.psych.60.110707.163612?journalCode=psych

McArdle, J. J. (2009). Latent variable modeling of differences and changes with longitudinal data. Annual Review of Psychology, 60(October 2008), 577–605. http://doi.org/10.1146/annurev.psych.60.110707.163612

McArdle, J. J., & Grimm, K. J. (2010). Five Steps in Latent Curve and Latent Change Score Modeling with Longitudinal Data. In Longitudinal Research with Latent Variables (pp. 245–273). Berlin, Heidelberg: Springer Berlin Heidelberg. http://doi.org/10.1007/978-3-642-11760-2_8

McArdle, J. J., & Hamagami, F. (2001a). Advanced Studies of Individual Differences Linear Dynamic Models for Longitudinal Data Analysis. In G. A. Marcoulides & R. E. Schumacker (Eds.), New Developments and Techniques in Structural Equation Modeling (pp. 203–246). London: Lawrence Erlbaum Associates Publishers. http://doi.org/10.1037/10409-005

McArdle, J. J., & Hamagami, F. (2001b). Latent difference score structural models for linear dynamic analyses with incomplete longitudinal data. In New methods for the analysis of change. Decade of behavior. (pp. 139–175).

McArdle, J. J., Hamgami, F., Jones, K., Jolesz, F., Kikinis, R., Spiro, A., & Albert, M. S. (2004). Structural modeling of dynamic changes in memory and brain structure using longitudinal data from the normative aging study. The Journals of Gerontology. Series B, Psychological Sciences and Social Sciences, 59(6), P294–304. http://doi.org/10.1093/GERONB/59.6.P294

McArdle, J. J., & Nesselroade, J. R. (1994). Using multivariate data to structure developmental change. Lawrence Erlbaum Associates, Inc.

McNeish, D., An, J., & Hancock, G. R. (2017). The Thorny Relation Between Measurement Quality and Fit Index Cutoffs in Latent Variable Models. Journal of Personality Assessment, 1–10. http://doi.org/10.1080/00223891.2017.1281286

McNeish, D., & Matta, T. (2017). Differentiating Between Mixed Effects and Latent Curve Approaches to Growth Modeling. Behavior Research Methods, (March). http://doi.org/10.13140/RG.2.2.21184.74243

Mehta, P. D., & West, S. G. (2000). Putting the individual back into individual growth curves. Psychological Methods, 5(1), 23–43. http://doi.org/10.1037/1082-989X.5.1.23

Meredith, W. (1993). Measurement invariance, factor analysis and factorial invariance. Psychometrika, 58(4), 525–543. http://doi.org/10.1007/BF02294825

Merkle, E. C., & Rosseel, Y. (2015). blavaan: Bayesian structural equation models via parameter expansion. arXiv, (Rosseel 2012). Retrieved from http://arxiv.org/abs/1511.05604

Merkle, E. C., You, D., & Preacher, K. J. (2016). Testing nonnested structural equation models. Psychological Methods, 21(2), 151–163. http://doi.org/10.1037/met0000038

Mills, K. L., Goddings, A.-L., Clasen, L. S., Giedd, J. N., & Blakemore, S.-J. (2014). The developmental mismatch in structural brain maturation during adolescence. Developmental Neuroscience, 36(3-4), 147–60. http://doi.org/10.1159/000362328

Millsap, R. E. (2011). Statistical approaches to measurement invariance. Routledge.

Moshagen, M., & Erdfelder, E. (2016). A New Strategy for Testing Structural Equation Models. Structural Equation Modeling: A Multidisciplinary Journal, 23(1), 54–60. http://doi.org/10.1080/10705511.2014.950896

Muetzel, R. L., Blanken, L. M. E., van der Ende, J., El Marroun, H., Shaw, P., Sudre, G., … White, T. (2017). Tracking Brain Development and Dimensional Psychiatric Symptoms in Children: A Longitudinal Population-Based Neuroimaging Study. American Journal of Psychiatry, appi.ajp.2017.1. http://doi.org/10.1176/appi.ajp.2017.16070813

Muthén, B., & Asparouhov, T. (2012). Bayesian structural equation modeling: A more flexible representation of substantive theory. Psychological Methods, 17(3), 313–335. http://doi.org/10.1037/a0026802

Muthén, L. K., & Muthén, B. O. (2002). How to Use a Monte Carlo Study to Decide on Sample Size and Determine Power. Structural Equation Modeling: A Multidisciplinary Journal, 9(4), 599–620. http://doi.org/10.1207/S15328007SEM0904_8

Muthén, L. K., & Muthén, B. O. (2005). Mplus: Statistical analysis with latent variables: User's guide. Los Angeles: Muthén & Muthén.

Neale, M. C. (2000). Individual fit, heterogeneity, and missing data in multigroup structural equation modeling. Lawrence Erlbaum Associates Publishers.

Newsom, J. T. (2015). Longitudinal structural equation modeling: A comprehensive introduction. London: Routledge.

Newton-Smith, W., & Lukes, S. (1978). The Underdetermination of Theory by Data. Proceedings of the Aristotelian Society, Supplementary Volumes, 52, 71–91-107. http://doi.org/10.2307/4106790

Oertzen, T., Hertzog, C., Lindenberger, U., & Ghisletta, P. (2010). The effect of multiple indicators on the power to detect inter-individual differences in change. British Journal of Mathematical and Statistical Psychology, 63(3), 627–646. http://doi.org/10.1348/000711010X486633

Olsson, U. H., Foss, T., Troye, S. V., & Howell, R. D. (2000). The Performance of ML, GLS, and WLS Estimation in Structural Equation Modeling Under Conditions of Misspecification and Nonnormality. Structural Equation Modeling: A Multidisciplinary Journal, 7(4), 557–595. http://doi.org/10.1207/S15328007SEM0704_3

Pearl, J. (2000). Causality: models, reasoning and inference. Cambridge: MIT press.

Pearl, J. (2012). The Causal Foundations of Structural Equation Modeling. Retrieved from http://ftp.cs.ucla.edu/pub/stat_ser/r370.pdf

Petscher, Y., Quinn, J. M., & Wagner, R. K. (2016). Modeling the co-development of correlated processes with longitudinal and cross-construct effects. Developmental Psychology, 52(11), 1690–1704.

Poldrack, R. A., & Gorgolewski, K. J. (2014). Making big data open: data sharing in neuroimaging. Nature Neuroscience, 17(11), 1510–1517. http://doi.org/10.1038/nn.3818

Quinn, J. M., Wagner, R. K., Petscher, Y., & Lopez, D. (2015). Developmental relations between vocabulary knowledge and reading comprehension: a latent change score modeling study. Child Development, 86(1), 159–75. http://doi.org/10.1111/cdev.12292

R Development Core Team. (2016). R: a language and environment for statistical computing. Vienna. Retrieved from http://www.r-project.org

Rajapakse, J. C., Giedd, J. N., & Rapoport, J. L. (1997). Statistical approach to segmentation of single-channel cerebral MR images. IEEE Transactions on Medical Imaging, 16(2), 176–186. http://doi.org/10.1109/42.563663

Rast, P., & Hofer, S. M. (2014). Longitudinal design considerations to optimize power to detect variances and covariances among rates of change: simulation results based on actual longitudinal studies. Psychological Methods, 19(1), 133–54. http://doi.org/10.1037/a0034524

Raykov, T., & Penev, S. (1999). On Structural Equation Model Equivalence. Multivariate Behavioral Research, 34(2), 199–244. http://doi.org/10.1207/S15327906Mb340204

Raz, N., Lindenberger, U., Rodrigue, K. M., Kennedy, K. M., Head, D., Williamson, A., … Acker, J. D. (2005). Regional brain changes in aging healthy adults: general trends, individual differences and modifiers. Cerebral Cortex (New York, N.Y.: 1991), 15(11), 1676–89. http://doi.org/10.1093/cercor/bhi044

Raz, N., Schmiedek, F., Rodrigue, K. M., Kennedy, K. M., Lindenberger, U., & Lövdén, M. (2013). Differential brain shrinkage over 6months shows limited association with cognitive practice. Brain and Cognition, 82(2), 171–180. http://doi.org/10.1016/j.bandc.2013.04.002

Ritchie, S. J., Bastin, M. E., Tucker-Drob, E. M., Maniega, S. M., Engelhardt, L. E., Cox, S. R., … Deary, I. J. (2015). Coupled Changes in Brain White Matter Microstructure and Fluid Intelligence in Later Life. Journal of Neuroscience, 35(22), 8672–8682. http://doi.org/10.1523/JNEUROSCI.0862-15.2015

Rodgers, J. L., & Lee, J. (2010). The epistemology of mathematical and statistical modeling: A quiet methodological revolution. American Psychologist, 65(1), 1–12. http://doi.org/10.1037/a0018326

Rosseel, Y. (2012). lavaan: An R Package for Structural Equation Modeling. Journal of Statistical Software, 10(2), 1–36.

Rosseel, Y. (2013). Longitudinal structural equation modeling. New York, (April).

Rovine, M. J., & Molenaar, P. C. M. (2001). A structural equations modeling approach to the general linear mixed model. In New methods for the analysis of change. (pp. 67–98). Washington: American Psychological Association. http://doi.org/10.1037/10409-003

Salthouse, T. A. (2014). Why are there different age relations in cross-sectional and longitudinal comparisons of cognitive functioning? Current Directions in Psychological Science, 23(4), 252–256. http://doi.org/10.1177/0963721414535212

Satorra, A., & Saris, W. E. (1985). Power of the likelihood ratio test in covariance structure analysis. Psychometrika, 50(1), 83–90. http://doi.org/10.1007/BF02294150

Schermelleh-Engel, K., Moosbrugger, H., & Müller, H. (2003). Evaluating the Fit of Structural Equation Models: Tests of Significance and Descriptive Goodness-of-Fit Measures. Methods of Psychological Research - Online, 8(2), 23–74.

Schmiedek, F., Lövdén, M., & Lindenberger, U. (2010). Hundred Days of Cognitive Training Enhance Broad Cognitive Abilities in Adulthood: Findings from the COGITO Study. Frontiers in Aging Neuroscience, 2(July), 1–10. http://doi.org/10.3389/fnagi.2010.00027

Schmiedek, F., Lövdén, M., & Lindenberger, U. (2014). Younger adults show long-term effects of cognitive training on broad cognitive abilities over 2 years. Developmental Psychology, 50(9), 2304–2310. http://doi.org/10.1037/a0037388

Segalowitz, S. J., & Rose-Krasnor, L. (1992). The construct of brain maturation in theories of child development. Brain and Cognition, 20(1), 1–7. http://doi.org/10.1016/0278-2626(92)90058-T

Sliwinski, M., Hoffman, L., & Hofer, S. M. (2010). Evaluating Convergence of Within-Person Change and Between-Person Age Differences in Age-Heterogeneous Longitudinal Studies. Research in Human Development, 7(1), 45–60. http://doi.org/10.1080/15427600903578169

Snitz, B. E., Small, B. J., Wang, T., Chang, C.-C. H., Hughes, T. F., Ganguli, M., … KLIEGEL, M. (2015). Do Subjective Memory Complaints Lead or Follow Objective Cognitive Change? A Five-Year Population Study of Temporal Influence. Journal of the International Neuropsychological Society, 21(9), 732–742. http://doi.org/10.1017/S1355617715000922

Steiger, J. H. (2007). Understanding the limitations of global fit assessment in structural equation modeling. Personality and Individual Differences, 42(5), 893–898. http://doi.org/10.1016/j.paid.2006.09.017

Steinberg, L. (2008). A Social Neuroscience Perspective on Adolescent Risk-Taking. Developmental Review: DR, 28(1), 78–106. http://doi.org/10.1016/j.dr.2007.08.002

Steyer, R., Schmitt, M., & Eid, M. (1999). Latent state-trait theory and research in personality and individual differences. European Journal of Personality, 13(5), 389–408. http://doi.org/10.1002/(SICI)1099-0984(199909/10)13:5<389::AID-PER361>3.0.CO;2-A

Stoel, R. D., Garre, F. G., Dolan, C., & van den Wittenboer, G. (2006). On the likelihood ratio test in structural equation modeling when parameters are subject to boundary constraints. Psychological Methods, 11(4), 439–455. http://doi.org/10.1037/1082-989X.11.4.439

Tohka, J., Zijdenbos, A., & Evans, A. (2004). Fast and robust parameter estimation for statistical partial volume models in brain MRI. NeuroImage, 23(1), 84–97. http://doi.org/10.1016/j.neuroimage.2004.05.007

Tomarken, A. J., & Waller, N. G. (2005). Structural equation modeling: strengths, limitations, and misconceptions. Annual Review of Clinical Psychology, 1, 31–65. http://doi.org/10.1146/annurev.clinpsy.1.102803.144239

Usami, S., Hayes, T., & McArdle, J. J. (2016). Inferring Longitudinal Relationships Between Variables: Model Selection Between the Latent Change Score and Autoregressive Cross-Lagged Factor Models Mail: Mail: Mail: University of S. Structural Equation Modeling: A Multidisciplinary Journal, 331–342.

van den Bos, W., & Eppinger, B. (2016). Developing developmental cognitive neuroscience: From agenda setting to hypothesis testing. Developmental Cognitive Neuroscience. http://doi.org/10.1016/j.dcn.2015.12.011

van der Sluis, S., Verhage, M., Posthuma, D., & Dolan, C. V. (2010). Phenotypic complexity, measurement bias, and poor phenotypic resolution contribute to the missing heritability problem in genetic association studies. PloS One, 5(11), e13929. http://doi.org/10.1371/journal.pone.0013929

Van Erp, S., Mulder, J., & Oberski, D. (2017). Prior sensitivity analysis in default Bayesian structural equation modeling. Psychological Methods. http://doi.org/10.17605/OSF.IO/5J3M9

van Harmelen, A.-L., Gibson, J. L., St Clair, M. C., Owens, M., Brodbeck, J., Dunn, V., … Goodyer, I. M. (2016). Friendships and Family Support Reduce Subsequent Depressive Symptoms in At-Risk Adolescents. PloS One, 11(5), e0153715. http://doi.org/10.1371/journal.pone.0153715

Vandenberg, R. J., & Lance, C. E. (2000). A Review and Synthesis of the Measurement Invariance Literature: Suggestions, Practices, and Recommendations for Organizational Research. Organizational Research Methods, 3(1), 4–70. http://doi.org/10.1177/109442810031002

Voelkle, M. C. (2007). Latent growth curve modeling as an integrative approach to the analysis of change. Psychology Science, 49(4), 375–414.

Voelkle, M. C., & Oud, J. H. L. (2015). Relating Latent Change Score and Continuous Time Models Relating Latent Change Score and Continuous Time Models. Structural Equation Modeling: A Multidisciplinary Journal, 22(3), 366–381. http://doi.org/10.1080/10705511.2014.935918

von Hippel, P. T. (2016). New Confidence Intervals and Bias Comparisons Show That Maximum Likelihood Can Beat Multiple Imputation in Small Samples. Structural Equation Modeling: A Multidisciplinary Journal, 23(3), 422–437. http://doi.org/10.1080/10705511.2015.1047931

von Oertzen, T., & Brandmaier, A. M. (2013). Optimal study design with identical power: An application of power equivalence to latent growth curve models. Psychology and Aging, 28(2), 414–428. http://doi.org/10.1037/a0031844

von Oertzen, T., Brandmaier, A. M., & Tsang, S. (2015). Structural Equation Modeling With Ωnyx. Structural Equation Modeling: A Multidisciplinary Journal, 22(1), 148–161. http://doi.org/10.1080/10705511.2014.935842

Wagenmakers, E.-J., & Farrell, S. (2004). AIC model selection using Akaike weights. Psychonomic Bulletin & Review, 11(1), 192–196. http://doi.org/10.3758/BF03206482

Weiskopf, N., Suckling, J., Williams, G., Correia, M. M., Inkster, B., Tait, R., … Lutti, A. (2013). Quantitative multi-parameter mapping of R1, PD^*^, MT, and R2^*^ at 3T: a multi-center validation. Frontiers in Neuroscience, 7, 95. http://doi.org/10.3389/fnins.2013.00095

Wendelken, C., Ferrer, E., Ghetti, S., Bailey, S. K., Cutting, L., & Bunge, S. A. (2017). Frontoparietal Structural Connectivity in Childhood Predicts Development of Functional Connectivity and Reasoning Ability: A Large-Scale Longitudinal Investigation. The Journal of Neuroscience: The Official Journal of the Society for Neuroscience, 37(35), 8549–8558. http://doi.org/10.1523/JNEUROSCI.3726-16.2017

Westfall, J., & Yarkoni, T. (2016). Statistically Controlling for Confounding Constructs Is Harder than You Think. PloS One, 11(3), e0152719. http://doi.org/10.1371/journal.pone.0152719

Wheaton, B., Muthen, B., Alwin, D. F., & Summers, G. F. (1977). Assessing Reliability and Stability in Panel Models. Sociological Methodology, 8, 84. http://doi.org/10.2307/270754

Whitaker, K. J., Vértes, P. E., Romero-Garcia, R., Váša, F., Moutoussis, M., Prabhu, G., … Villis, L. (2016). Adolescence is associated with genomically patterned consolidation of the hubs of the human brain connectome. Proceedings of the National Academy of Sciences of the United States of America, 113(32), 9105–10. http://doi.org/10.1073/pnas.1601745113

Wicherts, J. M., Dolan, C. V., & Hessen, D. J. (2005). Stereotype Threat and Group Differences in Test Performance: A Question of Measurement Invariance. Journal of Personality and Social Psychology, 89(5), 696–716. http://doi.org/10.1037/0022-3514.89.5.696

Wicherts, J. M., Dolan, C. V., Hessen, D. J., Oosterveld, P., van Baal, G. C. M., Boomsma, D. I., & Span, M. M. (2004). Are intelligence tests measurement invariant over time? Investigating the nature of the Flynn effect. Intelligence, 32(5), 509–537. http://doi.org/10.1016/j.intell.2004.07.002

Widaman, K. F., Ferrer, E., & Conger, R. D. (2010). Factorial Invariance within Longitudinal Structural Equation Models: Measuring the Same Construct across Time. Child Development Perspectives, 4(1), 10–18. http://doi.org/10.1111/j.1750-8606.2009.00110.x

Willis, S. L., & Schaie, K. W. (1986). Practical Intelligence: Nature and Origins of Competence in the Everyday World. (R. J. Sternberg & R. K. Wagner, Eds.). New York: Cambridge University Press. Retrieved from http://books.google.com/books?hl=nl&lr=&id=-Cw7AAAAIAAJ&pgis=1

Wothke, W. (1993). Nonpositive definite matrices in structural modeling. In K. A. Bollen (Ed.), Testing structural equation models (pp. 256–93). Newbury Park, CA: Sage.

Wothke, W. (2000). Longitudinal and multigroup modeling with missing data. In T. D. Little, K. U. Schnabel, & J. Baumert (Eds.), Modeling longitudinal and multilevel data: Practical issues, applied approaches, and specific examples (pp. 219–240). Mahwah, NJ: Lawrence Erlbaum Associates, Inc.

Wright, S. (1920). The Relative Importance of Heredity and Environment in Determining the Piebald Pattern of Guinea-Pigs. Proceedings of the National Academy of Sciences of the United States of America, 6(6), 320–32. Retrieved from http://www.ncbi.nlm.nih.gov/pubmed/16576506

Zhang, Z., Hamagami, F., Grimm, K. J., & McArdle, J. J. (2015). Using R Package RAMpath for Tracing SEM Path Diagrams and Conducting Complex Longitudinal Data Analysis. Structural Equation Modeling: A Multidisciplinary Journal, 22(1), 132–147. http://doi.org/10.1080/10705511.2014.935257

Ziegler, G., Ridgway, G. R., Blakemore, S.-J., Ashburner, J., & Penny, W. (2017). Multivariate dynamical modelling of structural change during development. NeuroImage, 147, 746–762. http://doi.org/10.1016/j.neuroimage.2016.12.017

